# Metabolic wastes are extracellularly disposed by excretosomes, nanotubes and exophers in mouse HT22 cells through an autophagic vesicle clustering mechanism

**DOI:** 10.1101/699405

**Authors:** Hualin Fu, Jilong Li, Peng Du, Weilin Jin, Daxiang Cui

**Affiliations:** Institute of Nano Biomedicine and Engineering, Department of Instrument Science and Engineering, School of Electronic Information and Electrical Engineering, Shanghai Jiao Tong University, 800 Dongchuan Road, Shanghai, 200240, P. R. China; National Center for Translational Medicine, Shanghai Jiao Tong University, 800 Dongchuan Road, Shanghai, 200240, P. R. China; Department of Colorectal Surgery, Xinhua Hospital, School of Medicine, Shanghai Jiao Tong University, Shanghai, China

**Keywords:** metabolic waste, excretosome, nanotube, exopher, clustering, vacuole, excretion

## Abstract

Neurodegenerative diseases are characterized by metabolic waste accumulations in neuronal cells. However, little is known about the mechanism of metabolic waste disposal. We showed that there is a constitutive waste excretion by nanotubes and structures called excretosomes in mouse neuronal HT22 cells under normal conditions and an exopher-type waste disposal under stressed conditions. Excretosomes and exophers are waste-enriched multivesicular bodies that are formed by autophagic vesicle clustering and fusions. Upon H_2_O_2_ treatment, metabolic waste marker intensified 1.4 to 3.4 fold. However, in dying H_2_O_2_-treated cells, glycolysis and stress response markers, alpha-Tubulin, Amyloid beta and alpha-Synuclein intensities reduced 30% to 80%. Amyloid beta, TAU and alpha-Synuclein associated with the autophagic vesicle clustering and fusion process and were excreted in large exophers by live oxidative-stressed cells before cell death with important implications for amyloid plaque, ghost tangle and Lewy body formation. H_2_O_2_ treatment also induced exophers containing nucleic acid materials and significantly increased the level of oxidative damage marker 8-OHdG. Oxidative stress-induced HT22 cell death could be due to the combinations of excretion failure, cytoskeleton disruption, defective glycolysis and DNA damage.

## Main Text

Neurodegenerative diseases with devastating symptoms such as dementia are affecting around 10 millions of people in China alone and 40 millions of people worldwide, which are serious threats to human health^1, 2^. Proteostatic imbalance accompanying with toxic protein aggregations such as Amyloid beta (Aβ), TAU, α-Synuclein (αSyn) or toxic protein modifications such as protein nitration, MDA modification, 4HNE modification is considered as a very important cause for neurodegenerative diseases such as Alzheimer’s disease ^3-6^ and Parkinson’s disease^7^. Metabolically damaged proteins and organelles are often marked with ubiquitin or LC3 for proteasome enzymatic degradation or autophagy-lysosome vesicle-type degradation^8, 9^. Oxidative stress increases the production of harmful metabolic waste products such as nitrotyrosine (nitroY), MDA or 4HNE-modified proteins^10-12^ and decreases proteasome, lysosome degradative functions. The cellular protein homeostasis appears to be largely balanced with synthesis and degradation. However, theoretically, the accumulated metabolites waste in cells could be disposed simply in a “throw away” process without going through the proteasome or lysosome degradation. On the other hand, not all of the metabolic wastes could be completely degraded into simple harmless building blocks and recycled, so that a “throw away” option should be always remained open. At the evolution level, it is well known that advanced organisms develop sophisticated excretion system to dispose waste materials. At the cellular level, accumulating evidence suggests that cells can also excrete waste or toxic substances. One example is the export of chemotherapy drugs by cancer cells either through multidrug resistance protein transporters or through exosomes ^13, 14^. Another example is the extracellular delivery of unneeded proteins by exosomes in the reticulocyte differentiation and maturation process^15^. Additionally, an important form of cellular waste or toxic material disposal is observed in the tunneling nanotubes, which can deliver damaged organelles such as mitochondria for transcellular degradation and chemotherapy drugs to connecting cancer cells^16,17^. Furthermore, a recent elegant C. elegans study discovered that the nematode neurons can dispose metabolic waste such as toxic protein aggregates and damaged mitochondria in large membranous structures called exophers^18^. These studies highlighted that cells can dispose toxic metabolites in a variety of ways; however, an integrated view of cellular waste excretion is still lacking.

Since we hypothesized that there exists a general waste excretion system at the cellular level, we decided to examine multiple metabolic waste marker behaviors in a simple mouse HT22 neuronal cell culture model. We then associated these metabolic waste markers with known components in regulating metabolic waste such as the autophagy-lysosome pathway, the ubiquitin pathway and the MVB (multivesicular body) pathway. The evidence we provided suggested that the cells can excrete condensed metabolic waste constitutively in particulated forms that we called excretosomes through cellular nanotubes under normal growth conditions with basal levels of oxidative stress. Additionally, under elevated oxidative stress conditions, the cells can dispose concentrated metabolic waste through large exophers. Our data indicated that excretosomes and exophers are formed by a waste condensation mechanism through autophagic vesicle clustering and fusion. Additionally, we showed that hydrogen peroxide oxidative stress induces cell death with the accumulation of metabolic wastes and DNA damage, the disruption of cytoskeleton, the depletion of vital glycolytic and stress response proteins but not the accumulation of Aβ, phos-TAU or αSyn. Our study also indicated that the autophagy-lysosome pathway, ubiquitin pathway and the MVB pathway converge on the regulation of cellular waste excretion.

## Results

### Constitutive excretion of metabolic waste in normally growing HT22 cells

To look for the existence of a cellular waste excretion system, we first searched the literature for general metabolic waste markers. Six markers were chosen in the experiments, which include three well-known metabolic waste markers related to oxidative stress and neurodegeneration such as MDA, 4HNE, nitroY and three general markers of metabolic waste regulation such as ubiquitin B (UBB), LC3 and LAMP2, aiming at the ubiquitin pathway, the autophagy pathway and lysosomes respectively. First, cultured mouse neuronal cell line HT22 cells on coverslips were stained by these six markers. We found that metabolic waste-related markers were often distributed at distant foci in particle-like forms away from cell-bodies under normal culture conditions (Fig. 1A). To emphasize their functions in metabolic waste excretion, we tentatively name these particles as excretosomes, which could be multi-vesicle membranous bodies containing heterogeneous waste materials, as shown more clearly in the following studies. Further we showed that many of these seemly randomly distributed excretosomes often associated with cellular nanotubes, especially at the terminals of nanotubes (Fig. 1B). Occasionally, some excretosomes presented as small plasma membrane blebbings close to cell bodies as illustrated in Supplementary Fig. 1. We believed that the same biochemical reactions were happening regardless excretosomes came out from nanotube terminal membrane puffs or membrane blebbings close to cell bodies. We also observed extracellular excretosomes that were not associated with cellular nanotube structures or plasma membranes (Fig. 1B, labeled with asterisks), which could be excretosomes excreted out from cells as shown later more clearly in Figure 6. In addition, we also noticed the extensive co-localization of LC3 with beta Actin (ACTB) and alpha-Tubulin (αTub) both in cellular nanotubes and in extracellular excretosomes (Fig. 1C). Furthermore, LC3 and mitochondria marker Fumarate Hydratase (FH) co-localized in the nanotubes or at extracellular locations indicating mitochondrial materials were also excreted. The brightly MDA antibody-stained excretosomes were usually small and had diameters ranged from 600 nm to 5 μm as measured by morphometry analysis with the Image J software. However, occasionally, larger excretosomes with weaker waste marker staining with the size up to 10 μm were also observed. To highlight the enrichment of waste markers in the excretosomes, we measured the signal intensities of the brightest excretosomes. They had on average 3 to 8 times higher intensity of metabolic waste marker staining comparing to average cell body staining with the respective value of MDA (4.31±0.31, p<0.001, degree of freedom (df) =38), 4HNE(3.83±0.58, p<0.001, df=35), nitroY (3.92±1.1, p<0.001, df=33), UBB (3.84±1.17, p<0.001, df=38), LC3 (7.69±1.59, p<0.001, df=38), and LAMP2 (4.14±1.52, p<0.001, df=38) (Fig. 1D). Although all six markers could be important indicators of cell excretion, given the abundant presence of LC3 and MDA in excretosomes, we chose LC3 and MDA as general cellular excretion markers and focused more on these two markers in the following studies.

**Fig. 1.**
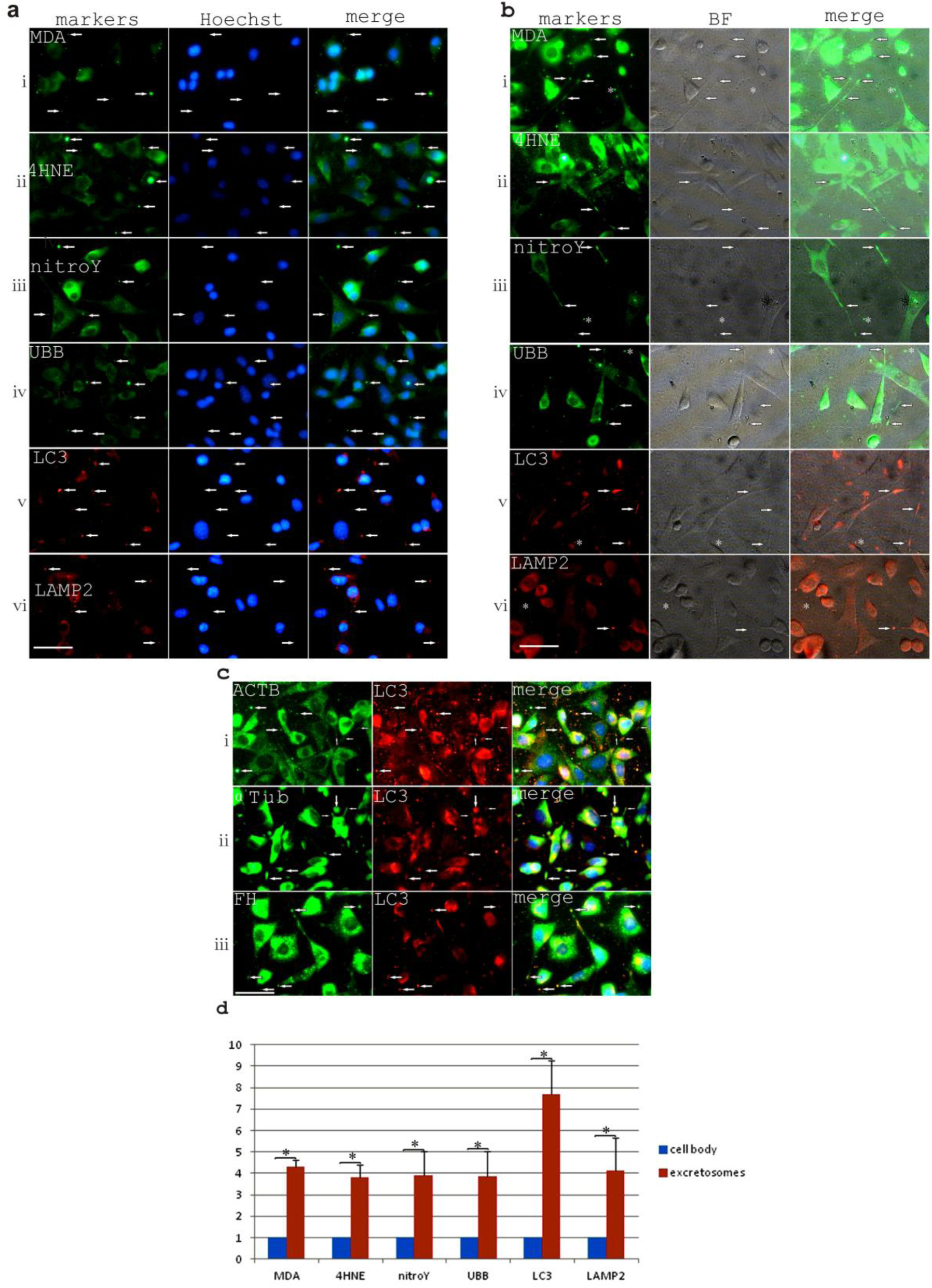
Metabolic wastes were disposed remotely away from cell bodies. **a.** Staining of MDA(i), 4HNE(ii), nitroY(iii), UBB(iv), LC3(v), LAMP2(vi) shows the frequent remote distribution of concentrated metabolic waste away from cell bodies, indicated with arrows. **b.** The merged fluorescence staining of MDA(i), 4HNE(ii), nitroY(iii), UBB(iv), LC3(v), LAMP2(vi) and bright field images showed that concentrated metabolic waste foci (excretosomes) were often associated with cellular nanotubes, either in the nanotubes or at the nanotube terminals, indicated by arrows. The asterisks showed some excretosomes that were not connected with the nanotubes, which could be excreted excretosomes. **c.** One of the metabolic waste excretion markers, LC3, co-localized with ACTB, αTub and mitochondria marker FH both in the cellular nanotube excretosomes and also extracellular excretosomes. Scale bars (**a-c**), 50 μm. **d.** The enrichment of metabolic waste markers in excretosomes compared to cell bodies, as measured by average area intensity.

### Exopher formation during hydrogen-peroxide oxidative stress treatment

During the course of H_2_O_2_ treatment, we also found that another form of waste disposal, the exopher pathway to export metabolic waste markers in large membranous structures is activated (Fig. 2a). At first, the cells form multiple membrane blebbing. Overtime, these membrane blebbings can develop into large exophers ranging from 2 to 20 μm in diameters. We proved that membrane blebbing and exophers are formed mostly by stressed but alive cells by Calcein AM and PI staining [Fig. 2b (i, ii)] although exopher formation can persist into some of the dying phase cells under continuous H_2_O_2_ treatment. In addition, we found that PI-negative HT22 cells contained high intensity of MDA in their exophers [Fig. 2a (i), 2b(iii)]. Exophers also enclosed mitochondrial materials marked with FH staining but with lower MitoTracker activity, indicating of the excretion of damaged mitochondria [Fig. 2b (iv)]. Exophers could be observed linking with the cell bodies by cellular nanotubes stained with GAPDH [Fig. 2b(v)] although standing-alone exophers with no nanotube attached were also observed at later stages of H_2_O_2_ treatment (Fig. 6b, Fig. 7b, d, f). Large exophers can form starting from small membrane blebbings within 10 minutes with continuous H_2_O_2_ treatment but other cells can have stabilized and unchanged membrane blebbings during the same time period of observation [Fig. 2b(vi)]. In the following studies, we considered large membrane blebbings during H_2_O_2_ treatment as early stage exophers. Excretosomes, membrane blebbings and late stage large exophers could all be clearly labeled by Calcein AM live imaging of H_2_O_2_ treated HT22 cells as indicated by Supplementary Fig. 1. Calcein AM dye is a membrane permeable fluorescent dye that can be cleaved off the acetoxymethyl (AM) ester group by non-specific esterases to yield bright green fluorescence. The cleaved fluorescent product became membrane impermeable. As shown in the following studies, excretosomes and exophers contain high amount of lipid vesicles that derived from ER or lysosomes, that contain high esterase activities. With reasons above, we believed that, Calcein AM labels some vesicles in the excretosome or exopher compartment. Metabolic wastes were more concentrated in blebbing stage exophers as we showed that metabolic waste markers intensified by 1.58 to 3.07 folds in the membrane blebbings, comparing to the rest of cell bodies with the respective value of MDA(1.93±0.41, p=0.008, df=18), 4HNE (1.58±0.3, p=0.008, df=18), nitroY (2.36±1.38, p=<0.001, df=34), UBB (2.26±0.72, p<0.001, df=20), LC3 (3.05±1.3, p=0.007, df=20), and LAMP2 (3.07±0.77, p<0.001, df=24) (Fig. 2c). When the late stage exophers grow larger, the exopher intensities of metabolic waste markers decrease.

**Fig. 2.**
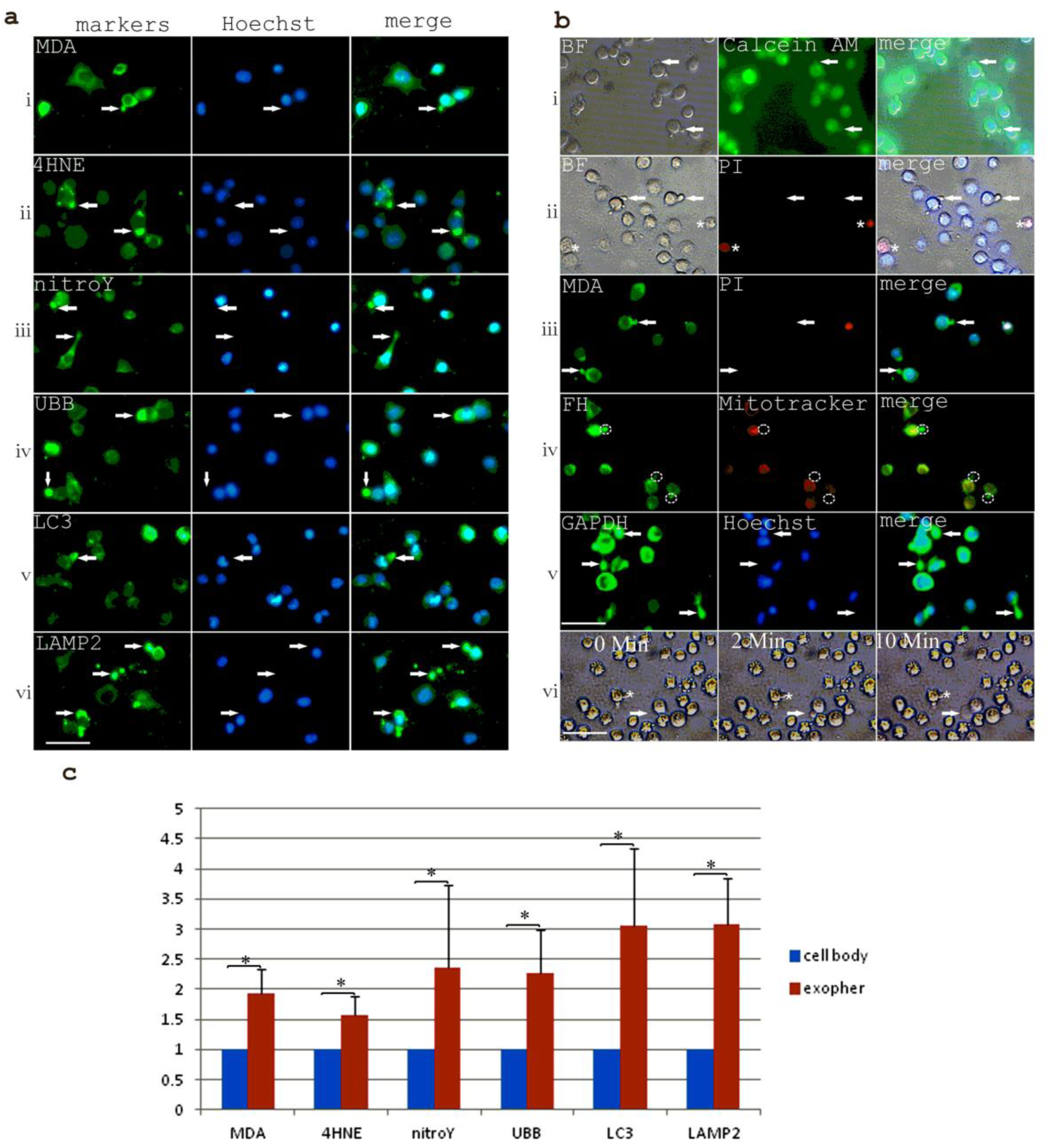
H_2_O_2_ treatment induced the export of metabolic waste through exophers. **a.** Membrane blebbing stage exophers were enriched for metabolic waste related markers such as MDA(i), 4HNE(ii), nitroY(iii), UBB(iv), LC3(v), LAMP2(vi), as indicated with arrows. Scale bar, 50 μm. **b.** Characterization of exophers during H_2_O_2_ treatment. **i.** Exopher-forming cells (indicated by arrows) were Calcein AM positive alive cells. **ii.** Exopher-forming cells (indicated by arrows) were PI-negative live cells. PI-positive dying cells were marked with “*”. **iii.** MDA-containing exopher formation, indicated by arrows, was happening in PI-negative cells. **iv.** Mitochondrial marker FH-stained exophers (indicated by circles) showed decreased MitoTracker staining. **v.** Some enlarged exophers (indicated by arrows and labeled with GAPDH antibody) could be seen linked with exopher-initiating cells by cellular nanotubes. **vi.** Real-time imaging showed that a large exopher (indicated by an arrow) could be formed starting from a small membrane blebbing within 10 minutes while another cell (indicated with “*”) showed unchanged membrane blebbing sizes during the same period of time. Scale bar in **a** and **b**(i-v), 50 μm. Scale bar in **b**(vi), 80 μm. **c**. Membrane protruding exophers contained concentrated metabolic waste related markers comparing to the rest of cell bodies when measuring the average area intensities.

### Oxidative stress induces cell death with accumulating metabolic wastes but not Aβ, αSyn, phos-TAU

Despite of the effort of metabolic waste excretion, under oxidative stress treatment with 1mM H_2_O_2_ for 4hrs, metabolic waste markers are accumulated significantly more in the treated HT22 cell bodies comparing to untreated cells (Fig. 3a) ranging from 1.37 fold to 3.41 folds with the respective value of MDA (2.87±1.04, p<0.001, df=54), 4HNE (2.53±0.81, p<0.001, df=38), nitroY (1.37±0.65, p=0.02, df=33), UBB (3.41±1.56, p<0.001, df=38), LC3 (3.08±1.97, p<0.001, df=38), and LAMP2 (1.41±0.46, P=0.002, df=38) (Fig. 3c). Further, it showed that metabolic waste markers are truly accumulated more in dying PI-positive H_2_O_2_-treated cells comparing to alive PI-negative H_2_O_2_-stressed cells(Fig. 3b) ranging from 1.4 folds to 4.3 folds when measuring the average area intensities with the respective value of MDA (1.42±0.33, p=0.004, df=25), 4HNE(1.42±0.49, p=0.003, df=18), nitroY (1.48±0.67, p=0.024, df=35), UBB (1.78±0.73, p=0.021, df=23), LC3 (4.25±2.41, p<0.001, df=25), and LAMP2 (1.93±0.82, p=0.0011, df=28) (Fig. 3d).

**Figure 3.**
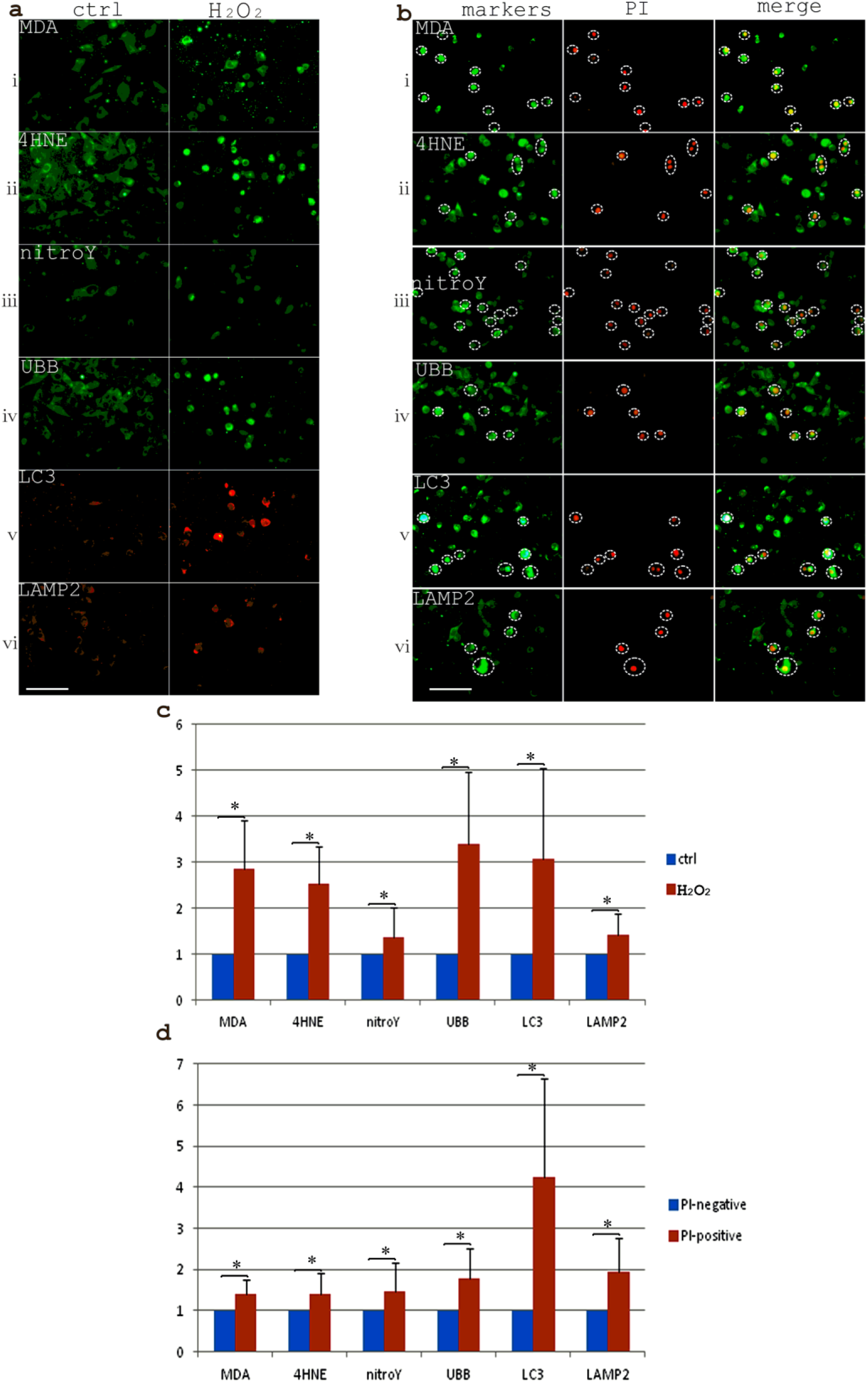
Metabolic wastes were accumulated in H_2_O_2_-treated cells and further enriched in dying cells. **a.** H_2_O_2_ treatment of HT22 cells. induced an accumulation of metabolic waste markers such as MDA(i), 4HNE(ii), nitroY(iii), UBB(iv), LC3(v) and LAMP2(vi). Scale bar, 100 μm. **b.** Most of the PI-positive dying HT22 cells (marked with circles) had strong metabolic waste related marker intensities comparing to PI-negative H_2_O_2_-stressed alive cells, indicated with arrows. The order of the markers was listed as MDA(i), 4HNE(ii), nitroY(iii), UBB(iv), LC3(v) and LAMP2(vi). Scale bar, 100 μm. **c.** The respective values for the enrichment of metabolic waste related markers in H_2_O_2_-treated HT22 cells vs. non-treated control HT22 cells. **d.** The respective values for the enrichment of metabolic waste related markers in H_2_O_2_-treated PI-positive dying HT22 cells comparing to H_2_O_2_-treated PI-negative alive HT22 cells.

Since Aβ, αSyn and phos-TAU were considered important metabolic stress signals inducing neuronal cell death, we studied these molecules in the same settings. We found that all these molecules are excreted constitutively in excretosomes under normal growth conditions (Fig. 4a-c). Additionally, αSYN and Aβ appeared to be associated with autophagic vesicle clustering and vacuole fusion and TAU could be also observed in large autophagic vacuoles. However, to our surprise, accumulation of Aβ, αSyn, phosphorylated-Tau was not observed in H_2_O_2_-treated HT22 cells (Fig. 4d, 4f). In fact, Aβ (0.30±0.11, p=0.003, df=35) and αSyn (0.46±0.09, p=0.02, df=37) intensities were reduced in dying PI-positive H_2_O_2_-treated HT22 cells compared to alive PI-negative H_2_O_2_-treated HT22 cells (Fig. 4e, 4g). We could observe some nuclear staining of these three markers in dying HT22 cells (Fig. 4e, Fig. 7d). Furthermore, analysis of glycolytic markers and stress-response proteins in the same settings also revealed that PKM2, GAPDH, ENO1, Hsp70, Annexin A2 (ANXA2), and G3BP1 had reduced intensities in dying PI-positive cells comparing to alive PI-negative H_2_O_2_-challenged HT22 cells with the respective value of PKM2 (0.16±0.04, p<0.001, df=18), GAPDH (0.43±0.13, p<0.001, df=26), ENO1 (0.32±0.11, p<0.001, df=49), Hsp70 (0.35±0.2, p<0.001, df=18), ANXA2 (0.65±0.28, p=0.02, df=18), G3BP1 (0.25±0.15, p<0.001, df=17) (Fig. 5a, 5d). These markers behave similarly as Aβ and αSyn, suggesting a defect of glycolysis and deficiency of stress responses might contribute to cell demise in this model. Moreover, exopher excretion of these markers was frequently observed, which potentially contribute to the rapid decrease of their protein expressions during H_2_O_2_ treatment (Supplementary Fig. 2). Since nanotubes and exophers are related to cytoskeleton networks, we also looked at the cytoskeleton changes with H_2_O_2_ treatment. Upon H_2_O_2_ treatment, αTub (0.29±0.14, p<0.001, df=17) but not beta Actin (ACTB) (1.06±0.35, p=0.58, df=40) intensity were greatly reduced in PI-positive cells compare to PI-negative H_2_O_2_-stressed cells (Fig. 5b, 5d). There was also a change of the spreading morphology of beta actin network to a round-up configuration in dying H_2_O_2_-treated cells, indicating that a reorganization of actin network was happening. Further, αTub loss was associated with LC3 accumulation in dying H_2_O_2_-treated HT22 cells suggesting microtubule defects associate with metabolic waste accumulation (Fig. 5c).

**Fig. 4:**
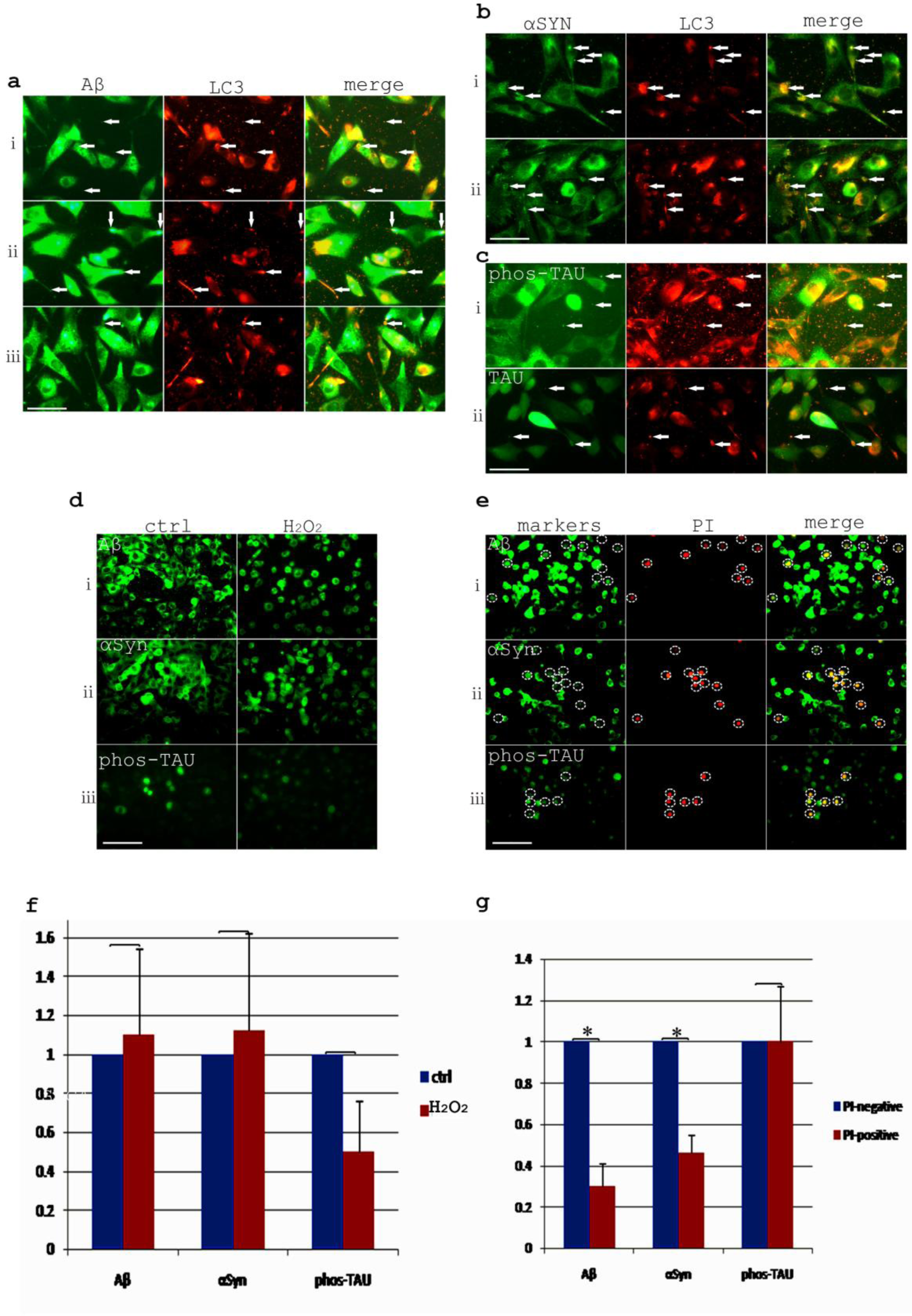
Aβ, αSYN, phos-Tau was not significantly accumulated in oxidative stressed dying HT22 cells. **a.** Aβ was associated with LC3 staining in excretosomes (i), clustered autophagic vesicles in nanotubes (ii) and in large autophagic vacuoles (iii). Arrows indicated the co-localizations. **b.** αSYN was associated with LC3-labeled excretosomes, intracellular clustered autophagic vesicles (i) and large autophagic vacuoles (ii). Arrows indicated the co-localizations. **c.** TAU was associated with LC3-labeled excretosomes (i) and phos-TAU co-localized with LC3-labeled large autophagic vacuoles (ii). Arrows indicated the co-localizations. Scale bar in a-c, 50 μm. **d.** Aβ, αSYN, phos-TAU was not significantly accumulated in H_2_O_2_-treated HT22 cells. **e.** Aβ, αSYN, phos-TAU was not significantly accumulated in dying H_2_O_2_-treated HT22 cells comparing to alive H_2_O_2_-treated HT22 cells. Circles indicated the PI-positive dying HT22 cells. Scale bar in d and e, 100 μm. **f.** quantitation of Aβ, αSYN, phos-TAU protein intensity in H_2_O_2_-treated cells comparing to untreated HT22 cells. **g.** quantitation of Aβ, αSYN, phos-TAU protein intensity in dying H_2_O_2_-treated cells comparing to alive H_2_O_2_-treated HT22 cells.

**Fig. 5.**
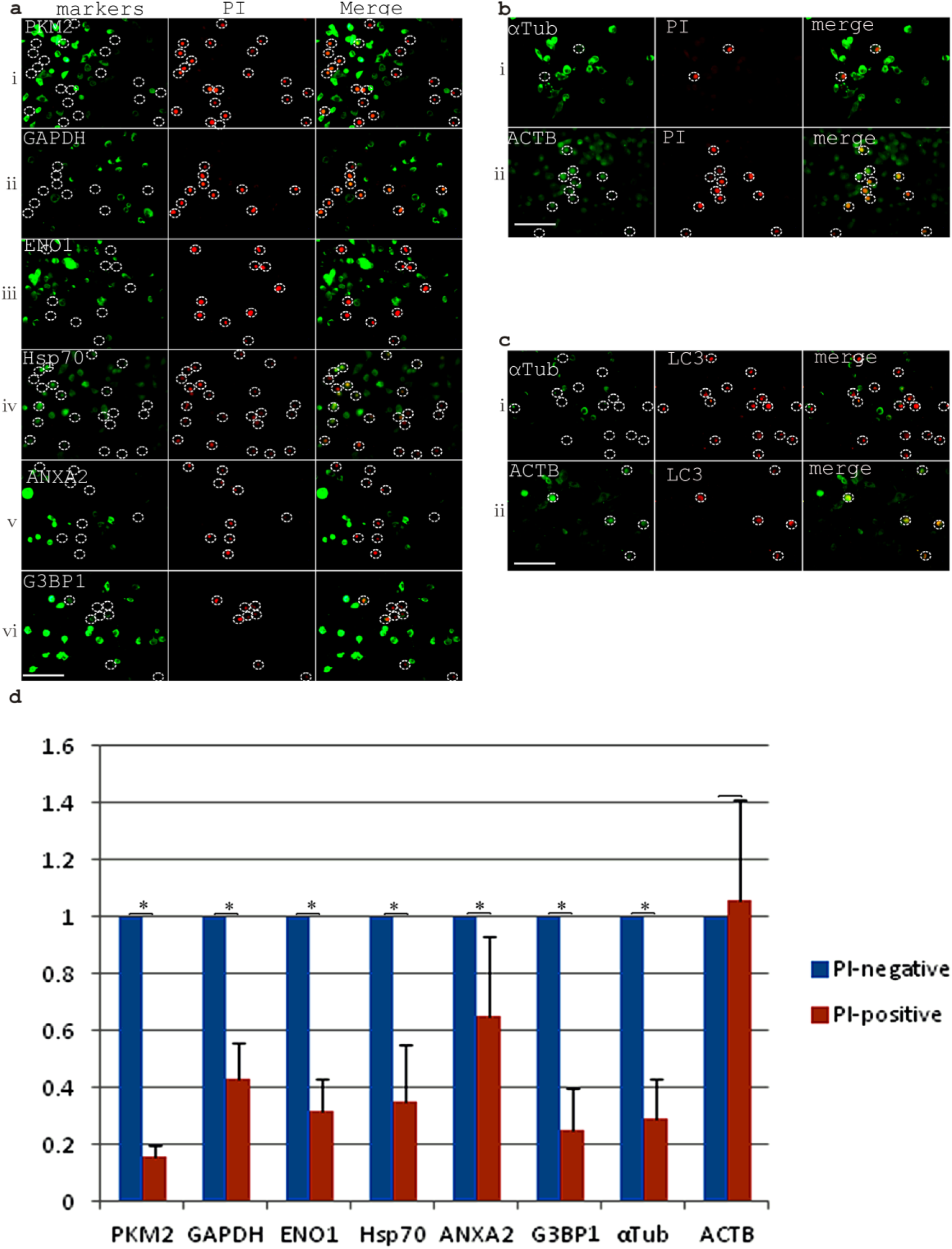
Analysis of glycolysis, stress-related markers and cytoskeleton proteins in PI-positive dying H_2_O_2_-treated cells. **a.** Decreased expression of PKM2(i), GAPDH(ii), ENO1(iii), HSP70(iv), ANXA2(v), G3BP1(vi) in PI-positive dying H_2_O_2_-treated cells comparing to PI-negative live H_2_O_2_-treated cells. Circles indicated the PI-positive dying HT22 cells. **b.** Decreased protein intensity of αTub (i) but not ACTB (ii) in PI-positive dying H_2_O_2_-treated cells comparing to PI-negative live H_2_O_2_-treated cells. Circles indicated the PI-positive dying HT22 cells. **c.** The disappearance of αTub (i) but not ACTB(ii) expression is coincident with the accumulation of LC3 staining. Circles indicated the LC3-accumulating HT22 cells. Scale bars (**a-c**), 100 μm. **d.** Quantitation showed that PKM2, GAPDH, ENO1, HSP70, ANXA2, G3BP1, αTub protein expression densities were reduced significantly in PI-positive dying H_2_O_2_-treated cells comparing to H_2_O_2_-treated PI-negative live cells.

### Excretosomes are formed by autophagic vesicle clustering and fusion

To understand how excretosomes are formed, we tracked the flow of metabolic waste materials labeled by LC3 from intracellular locations to extracellular locations. We found that autophagic vesicles were clustered inside of the normally growing HT22 cells. Particularly, autophagic vesicles can be seen clustered in puff-like cellular domains, in cellular nanotubes, and in an array of vesicle clusters in cellular nanotubes (Fig. 6a). In addition, a large autophagic vacuole formed from autophagic vesicle fusion could be frequently observed at the cellular terminals with the diameters around 5.7 um (5.70±1.01, N=7). Further, these autophagic vesicle clusters or fused vacuoles could be seen released from nanotube terminals to form extracellular excretosomes (Fig. 6b). The size and compactness of autophagic vesicle clusters varied between different excretosomes, which explains the heterogeneous appearance of excretosomes. The extracellular distribution of excretosomes, the often peripheral and distal autophagic vesicle clustering all suggested the possibility of a polarized transport of autophagic vesicles. We checked the co-staining of autophagic maker LC3 and microtubule marker alpha-Tubulin. The data showed abundant co-localization of LC3 and αTub in excretosomes, at intracellular vesicle clustering sites, in the cellular nanotubes and in the large autophagic vacuoles (Fig. 6c). The autophagic vesicle clustering and fusion process was possibly facilitated by microtubule directed transport. Clips from a real time recording of a Calcein AM-labeled vesicle away from cell body and magnified pictures of autophagic vesicle clustering and association with microtubules could be seen in Supplementary Fig. 3. Autophagic vesicle clustering and fusion likely simultaneously promote autophagic waste degradation and excretosome disposal. Since autophagic vesicles are thought to be derived from ER and ER is considered a major organelle containing internal calcium stores, we proposed autophagic vesicle clustering and fusion likely increased local calcium concentrations. A previous study already showed that increased calcium in the autophagic compartments upon autophagy induction in K562 cells^19^. We checked a marker of calcium signaling activation by phos-CAMK2 staining and found that excretosomes were positive for this marker. Additionally, phos-CAMK2 co-localized with autophagic vesicle clustering in the nanotubes and in the enlarged nanotube terminal vacuoles (Fig. 6d). Calcium signaling is possibly related to the autophagic vesicle clustering, fusion and disposal process. To address the regulatory factors involving autophagic vesicle clustering and fusion further, we checked the interaction of LC3 with an MVB marker CD63, which is also a lipid raft tetraspanin protein that could form membrane microdomains by mediating homotypical or heterotypical interactions. We found abundant co-localization of CD63 and LC3 in excretosomes, at intracellular autophagic vesicle clustering sites and in autophagic vesicle clustering in the cellular nanotubes and also in the enlarged autophagic vacuoles (Fig. 6e). Previous research indicated that autophagic vesicles could fuse with MVBs to form amphisomes^19^. Autophagic vesicle clustering likely involve the interactions between autophagosomes and MVBs and tetraspanin proteins such as CD63 possibly play important roles during this process. Higher magnification pictures of Calcein AM multivesicular labeling and CD63-LC3 co-immunostaining in Supplementary Fig. 4 illustrated the multivesicular feature of excretosomes in more detail. Furthermore, we also checked the co-localization of ubiquitin B and autophagic vesicles. UBB co-localized with LC3 signals in excretosomes, in intracellular condensations and autophagic vesicle clustering in the cellular nanotubes, and in the enlarged autophagic vacuoles (Fig. 6f). Ubiquitin pathway likely also participate the autophagic vesicle clustering and fusion process through ubiquitin-mediated protein interactions.

The condensations and excretions of other metabolic waste markers such as MDA, 4HNE, nitroY were also tightly linked to the autophagy pathway. We found MDA co-localized with LC3 in intracellular condensations, in the nanotubes and in the fused large autophagic vacuoles (Fig. 6g). The clustering of LC3 and MDA signals was also happening intracellularly and extracellularly and in exophers in H_2_O_2_-treated HT22 cells, suggesting similar mechanism is operating even when cells are facing increased oxidative stress (Fig. 6h). Calcein AM labeling of exophers with multivesicular features and enlarged pictures showing the clustering of autophagic vesicles in exophers and also the enlarged autophagic vacuole in membrane blebbings of H_2_O_2_-treated HT22 cells could be seen in Supplementary Fig. 5. The excretosomes and exophers are thus biochemically equivalent structures although at different scales. A similar co-localization of 4HNE and nitroY signals with autophagic vesicle clustering were shown in Supplementary Fig. 6 and Fig. 7. To further prove the pivotal role of autophagy in regulating metabolic waste excretion, we treated HT22 cells with both H_2_O_2_ and an autophagy inhibitor Chloroquine in a 3-hour experiment. The co-treatment of 1mM H_2_O_2_ and Chloroquine (100 μM) induced almost 100% cell death while treating with H_2_O_2_ alone for 3hrs induces only a few dying HT22 cells (Fig. 6i). Apparently, autophagy inhibition by Chloroquine accelerated excretion failure with increased MDA accumulation and increased cell death upon H_2_O_2_ treatment.

**Fig. 6.**
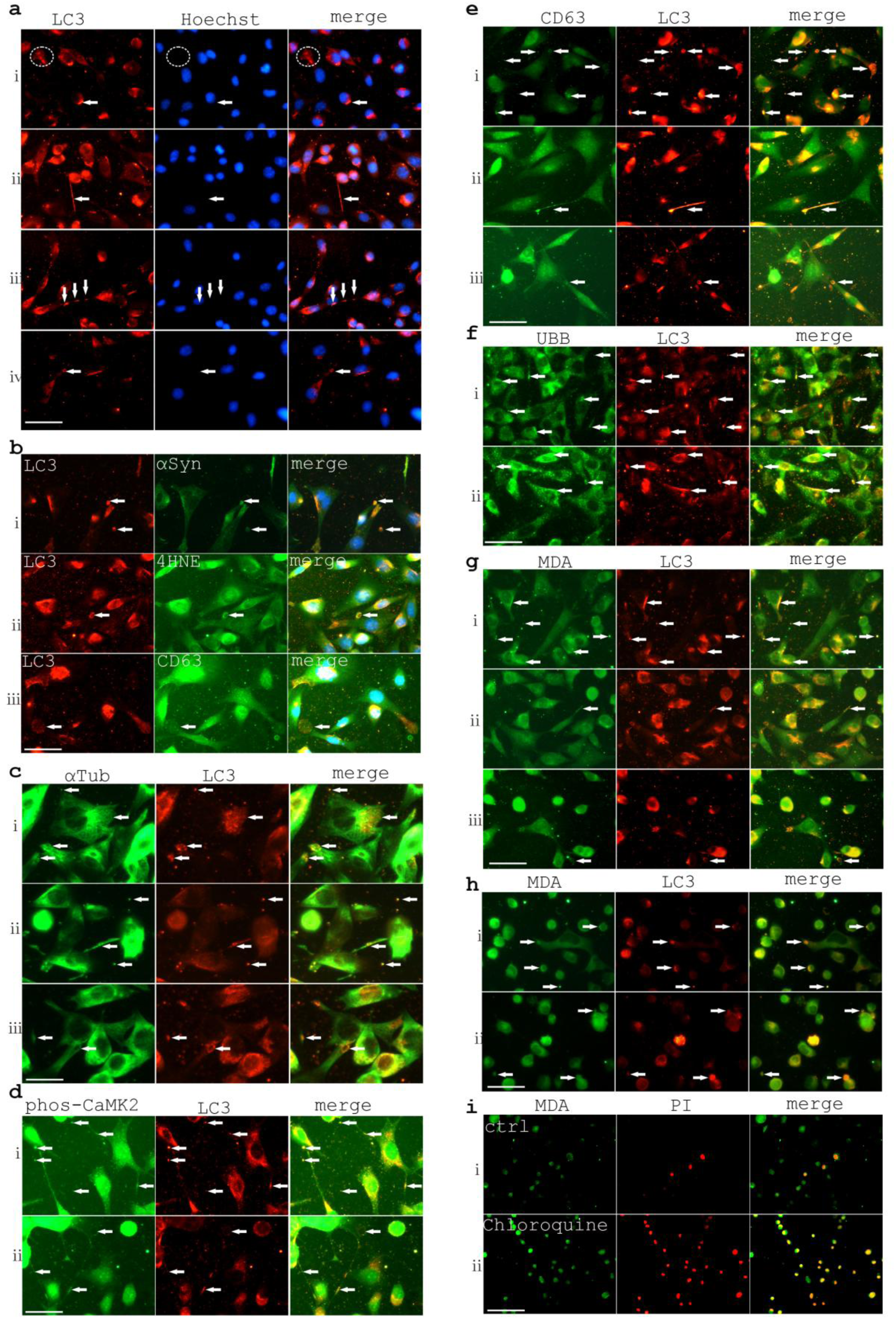
Excretosome formation and metabolic waste excretion are associated with autophagic vesicle clustering and fusion. **a.** Autophagic vesicle clustering and fusion in HT22 cells detected with LC3 immuostaining. **i.** Puff-like condensation of autophagic vesicles, indicated by a circle. **ii.** Vesicle clustering in cellular nanotubes (arrows). **iii.** An array of three auophagic vesicle clusters in the cellular nanotube (arrows). **iv.** An enlarged autophagic vacuole likely arising from vesicle fusion with an approximate size of 4 μm (indicated with an arrow). **b.** Autophagic clusters or vacuoles were excreted at cell terminals. **i.** Clustered autophagic vesicles were released at cell terminals with αSYN co-staining (arrows). **ii.** A large autophagic vacuole with 4HNE co-staining was released (arrow). **iii.** A large less-condensed vesicle cluster with clear characteristics of multivesicular bodies was released at cell terminals with CD63 costaining (arrow). **c.** Autophagic vesicles colocalized with microtubules labelled with αTub at intracellular vesicle clustering site and also in excretosomes (i) and in nanotube vesicle clusters (ii) and in enlarged nanotube terminal vacuoles (iii) indicated with arrows. **d.** phos-CAMK2 staining colocalized with LC3-labeled autophagic vesicle signals in excretosomes and in nanotubes (i) and in nanotubes and vacuoles (ii) indicated with arrows. **e.** Autophagic vesicle clustering and fusion was associated with CD63 expression in the same microdomains. CD63 colocalization with LC3 in excretosomes and intracelluar condensations (i) and nanotubes (ii) and in an enlarged CD63-LC3 double-positive vacuole possbily arising from autophagosome and MVB vesicle fusion (iii), indicated with arrows. **f.** UBB colocalized with LC3 in excretosomes, clustered autophagic vesicles (i) and enlarged nanotube terminal vacuoles(ii), indicated with arrows. **g.** In wild type HT22 cells, MDA colocalized with LC3 in intracellular condensations of autophagic vesicles (i) and in clustered autophagic vesicles in a cellular nanotube (ii) and in an enlarged autophagic vacuole possbily arising from vesicle fusion with an approximate size of 7 μm (iii), indicated with arrows. **h.** Intense autophagic vesilce clustering in H_2_O_2_-treated HT22 cells. MDA colocalized with clustered autophagic vesicles both intracellularly and extracellularly (i) and in exophers. and membrane blebbings (ii), indicated with arrows. **i.** Autophagy inhibitor Chloroquine treatment increased HT22 cell death with accelerated metabolic waste MDA accumulation. (i) MDA and PI staining of cells that were treated with 1mM H_2_O_2_ for 3 hrs. (ii) MDA and PI staining of cells that were treated with 1mM H_2_O_2_ for 3 hrs in the presence of 100 μM Choloroquine. Scale bars in **a**, **b**, **d**-**h**, 50 μm. Scale bar in **c**, 25 μm. Scale bar in **i**, 100 μm.

**Fig.7.**
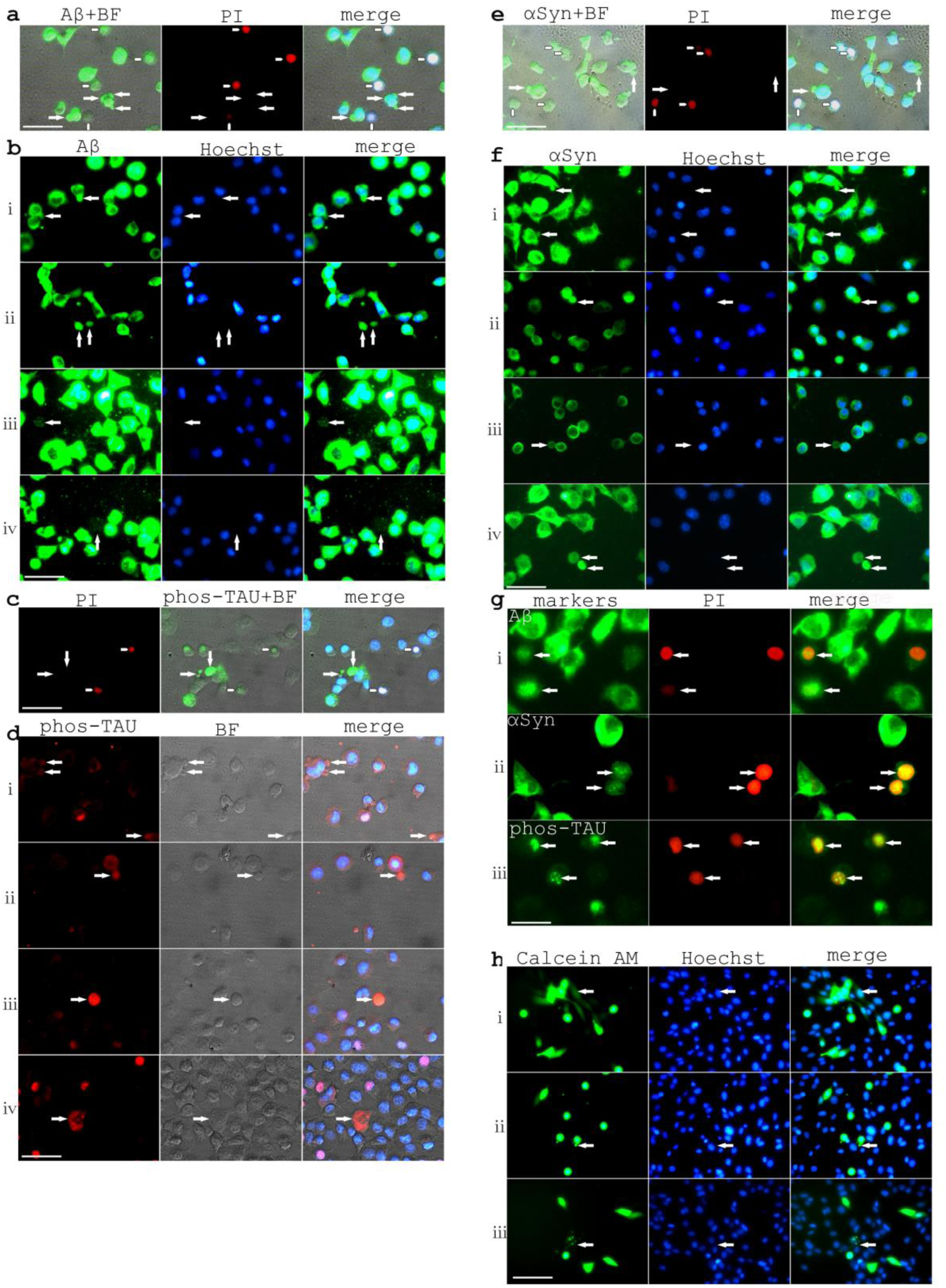
Extracellular deposits of Aβ, phos-TAU, and αSyn in exophers by live HT22 cells under stress. **a.** PI-negative HT22 cells showed Aβ-positive exopher (longer arrows) formation. The shorter arrows indicate PI-positive cells. **b.** Different stages of Aβ exopher and Aβ-positive plaque formation. The arrows indicate exophers or plaques. **c.** PI-negative HT22 cells showed phos-TAU-positive exopher (longer arrows) formation. The shorter arrows indicate PI-positive cells. **d.** Different stages of phos-TAU-positive exopher formation and extracellular deposits, indicated with arrows. **e.** PI-negative HT22 cells showed αSyn-positive (longer arrows) exopher formation. The shorter arrows indicate PI-positive cells. **f.** Different stages of αSyn-positive exopher formation and extracellular deposits, indicated with arrows. Scale bars, 50 μm. **g.** Nuclear localization of Aβ, αSyn, phos-TAU in dying HT22 cells. Scale bar, 25 μm. **h.** Possible modes of intercellular transmission in HT22 cells suggested by Calcein AM staining. (**i**). Possible transmission by cytoplasmic nanotubes showing by the diffusive cytoplasmic pattern of fluorescence in recipient cells. (**ii**). Calcein AM-positive donor cells with membrane blebbing formation. (**iii**). Recipient cells showed granule-like patterns, possibly received multiple vesicles from exophers. Scale bar, 100 μm.

### Aβ, phos-TAU and αSyn are extracellularly deposited by live cells under stress

When going through the data, it is to our surprise that Aβ, αSyn expression reduced in dying H_2_O_2_-treated cells, which seems to contradict with previous literature. In neurodegenerative diseases, the question still remains is whether the pathological deposits of Aβ, αSyn, and phos-TAU are primary causes of diseases or important secondary phenotypes. We provided evidence to show that Aβ, αSyn, and phos-TAU were excreted in large quantities in the form of exophers by live neuronal cells (Fig. 7). H_2_O_2_ treatment induced Aβ-positive exopher formation starting from membrane blebbings in PI-negative cells, developing into large exophers and diffusive plaque-like structures overtime. The diffusive Aβ staining plaques probably derived from broken and leaking Aβ exophers (Fig. 7a, b). Additionally, H_2_O_2_ stress induced phos-TAU-positive exopher formation in PI-negative cells, with many standing alone extracellular phos-TAU-positive deposits could be observed (Fig. 7c, d). Furthermore, H_2_O_2_ treatment stimulated αSyn-positive exopher formation starting from membrane blebbings in PI-negative cells, which then developed into large linking exophers or standing alone exophers (Fig. 7e, f). The data indicated that, in this cell line model, large extracellular deposits of Aβ, αSyn, and phos-TAU are important phenotypes during the severe oxidative-stress response. The extracellular deposits of Aβ, phos-TAU and αSyn in exophers under oxidative stress probably could explain the mechanism of the generation of amyloid plaques, ghost tangles in Alzheimer’s disease^20, 21^ and Lewy body formation in Parkinson’s disease^22^. Apparently, in dying neurons, Aβ and αSyn show a weak nuclear staining but phos-TAU stains strongly in the nuclei [Fig. 7g]. The nuclear functions of these proteins are not clear at present. We also showed that neurons cells probably transmit materials through both nanotubes and exophers by Calcein AM dye prelabeling and transmission experiment in HT22 cells [Fig. 7h]. The pathological phenotypes of Aβ, αSyn, phos-TAU in neurodegenerative disease possibly rise from oxidative stress and a failure of waste clearance.

### The convergence of autophagy, MVB, lysosome pathways on waste excretion

To further characterize autophagy-related signaling pathways in the cellular excretion system, more immunostaining experiments aiming at autophagy, MVB and lysosome pathways were performed. We found that MDA co-localized with autophagy marker LC3 and another MVB marker and also lipid raft protein CD9 and lysosome marker LAMP2 in excretosomes of normal growing cells, suggesting that auto-lysosomes and MVBs both play important roles of MDA excretion (Fig. 8a). In addition, we found that LC3, CD9 and LAMP2 vesicles clustered in the cellular nanotubes and the large nanotube terminal vacuoles were also positive for LC3, CD9 and LAMP2 (Fig. 8b, c), indicating a convergence of autophagy-lysosome and MVB pathway on cellular excretion. The autophagic vesicle clustering and fusion process is apparently happening together with lysosome and MVB clustering and fusion. Since cellular nanotube transport is linked with autophagic vesicle clustering and fusion process, it is important to look into the nanotube changes upon oxidative stress treatment. In Fig. 8d, it showed, by αSYN staining, H_2_O_2_ treated HT22 cells had reduced nanotube formation comparing to control untreated cells, which indicates that nanotube loss might contribute to the excretion failure phenotype. The nanotube loss phenotype could be due to the severe disruption of cytoskeleton structure during H_2_O_2_ treatment. As shown in Fig. 8e, exophers contained large amounts of αTub and ACTB. Additionally, H_2_O_2_ treatment induced actin ring formation at the bases of membrane blebs, suggesting a membrane damage and repair response was taking place during exopher disposal. It is known that oxidative stress not only induces proteostatic imbalance but also induces DNA damage, we checked the DNA damage markers during H_2_O_2_ treatment (Fig. 8f). At first sight, we noticed that DNA materials existed in exophers labeled by Calcein AM dye. Further, we found that phos-p53 positive micro nuclei DNA detached from the parent nuclei and also phos-p53 positive DNA existed in exophers extracellularly. Nuclear DNA excretion is likely linked to nuclear material autophagy, coined with a specific name of nucleophagy, as previously described^23, 24^. We also observed the distribution of oxidative nucleic acid damage marker 8-OHdG in the HT22 exophers, suggesting that damaged nucleic acid materials could also be excreted in exophers. Treatment of H_2_O_2_ induced roughly 5 times more intense nuclear 8-OHdG antibody staining in severely affected HT22 cells (4.90±0.74, p<0.001), suggesting extensive DNA damage is another factor contributing to H_2_O_2_-induced cell death [Fig. 8f(v)]. Furthermore, exophers induced by H_2_O_2_ treatment contained active Caspase-3, phos-p38 and phos-p53. In PI-positive dying HT22 cells, Caspase-3 activation and nuclear staining of phos-p38 and phos-p53 was also observed, indicating the possible involvement of Caspase-3, p38 and p53 in H_2_O_2_ induced cell death (Supplementary Fig. 8). Nuclear activation of phos-p53 could be due to increased oxidative DNA damage in H_2_O_2_-treated HT22 cells as suggested by 8-OHdG staining shown in Fig. 8f.

**Fig. 8.**
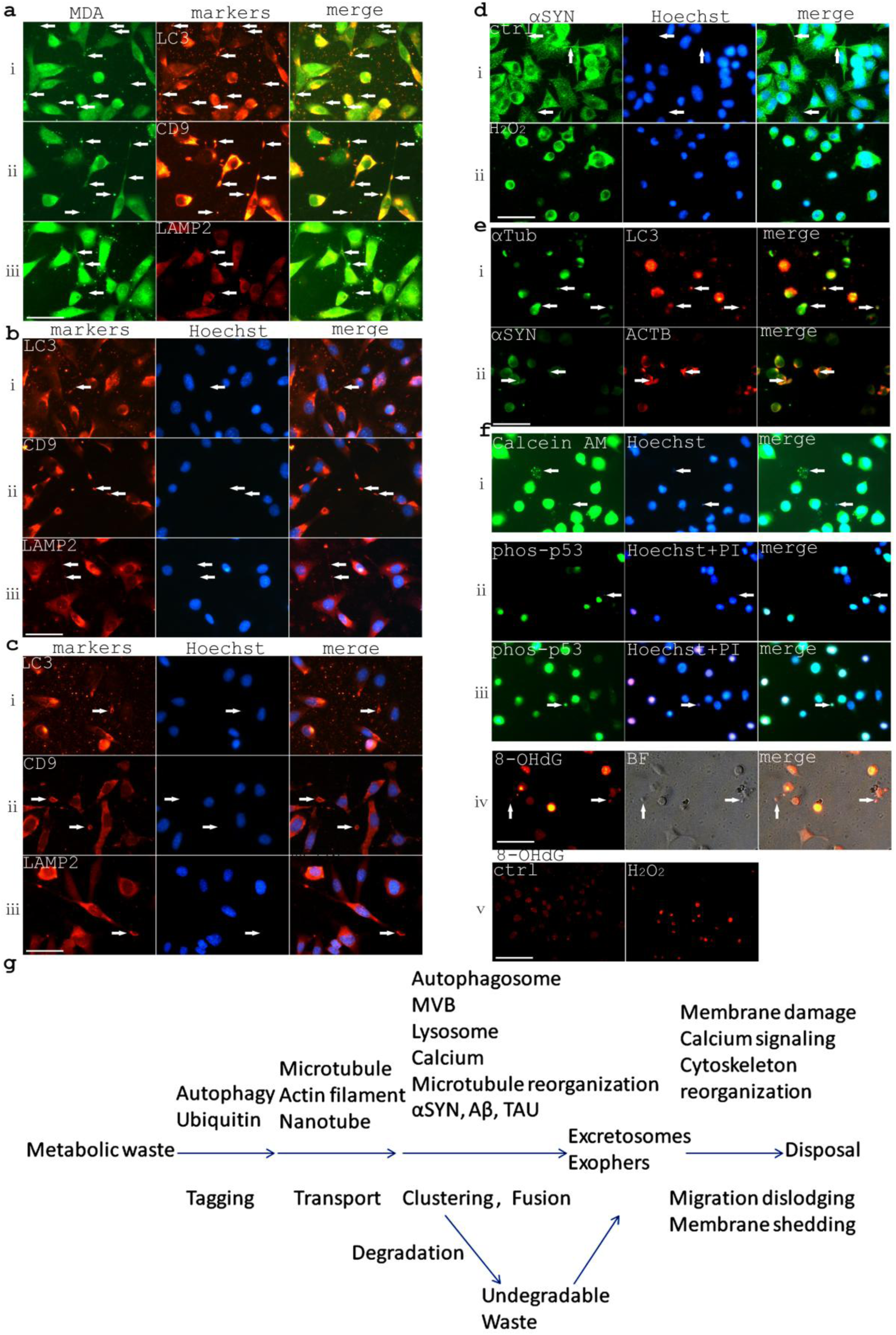
Autophagy, MVB and lysosome all contributed to metabolic waste removal through vesicle co-clustering and fusion. **a.** MDA signals co-localized with autophagy marker LC3, MVB marker CD9 and lysosome marker LAMP2. The arrows indicated the co-localizations. **b.** LC3, CD9 and LAMP2 vesicles clustered in nanotubes, indicated with arrows. **c.** LC3, CD9 and LAMP2 staining co-localized in cellular nanotube terminal vacuoles, indicated with arrows. **d.** Nanotube loss phenotype with H_2_O_2_ treatment as shown by αSYN immunostaining. **i.** Control. **ii.** H_2_O_2_-treated cells. Arrows indicated the nanotubes. **e.** Exopher formation in H_2_O_2_-treated HT22 cells carried away αTub (i) and ACTB (ii) and also induces actin ring-like structure formation at the base of membrane blebs (indicated with arrows). **f.** H_2_O_2_ treatment also increased DNA damage in HT22 cells significantly. **i.** Calcein AM labeled exophers contain DNA from H_2_O_2_-challenged HT22 cells, indicated with arrows. **ii.** phos-p53 labeling of detaching nuclear DNA in H_2_O_2_-treated HT22 cells, indicated with arrows. **iii.** phos-p53 labeling of exopher DNA from H_2_O_2_-treated HT22 cells, indicated with arrows. **iv.** Exophers and membrane blebbings contain 8-OHdG from H_2_O_2_-treated HT22 cells, indicated with arrows. **v.** Significantly elevated 8-OHdG staining upon H_2_O_2_ treatment of HT22 cells comparing to control untreated cells. Scale bars, 50 μm except the scale bar in **f**(iv) is 100 μm. **g.** A simplified scheme showing the framework of a cellular excretion system based on excretosomes and exophers.

To investigate whether excretosomes or exopher excretion is a general way of waste disposal upon stress in different cell types, we repeat some of the experiments done in HT22 cells in another cell line, the glioma cell line U251 cells. We noticed that U251 cells also disposed excretosomes with positive LC3 signals. Intracellular autophagic vesicle clustering was also observed in U251 cells (Supplementary Fig. 9). Live and stressed U251 cells also produced exophers (Supplementary Fig. 10). We additionally observed Aβ, αSyn, TAU, Cytochrome C (CytC), HMGb1, LC3, MDA expressed in U251 exophers upon H_2_O_2_ challenge (Supplementary Fig. 10, Fig. 11). By PI labeling of dying U251 cells, we confirmed that Aβ, αSyn, TAU were not elevated in dying U251 cells. Instead, decreased Aβ, αSyn staining in dying cells was observed (Supplementary Fig. 12). At the same time, the stress response protein G3BP1 also showed decreased staining in the dying U251 cells. Nuclear TAU, phos-p38 and p53 staining in the dying U251 cell nuclei suggested that TAU, phos-p38 and p53 signaling could be activated during oxidative-stress-induced U251 cell death similarly as shown in HT22 cells. Dye transmission experiment was also repeated in U251 cells, confirming that nanotubes and exophers could be two ways of material transfer between U251 cells (Supplementary Fig. 13).

Furthermore, to examine if excretosomes and exophers exist in human neuronal tissues *in vivo*, we performed immunostaining experiments on gastric myenteric neurons. First, we observed Aβ, αSyn and TAU staining in the gastric myenteric plexus by co-staining with a pan-neuronal marker UCHL1 (Supplementary Fig. 14). αSyn, TAU expressed strongly in both gastric neurons and glial cells while Aβ stained stronger in gastric neurons than associated glial cells. The co-staining of an astroglial cell marker S100 and UCHL1 showed the tight association of neurons and their accessory glial cells. The tight spacial wrapping of neurons by glial cells makes the specific detection of excretosomes and exophers from gastric neurons difficult. However, we were able to detect focal expression of MDA co-localizing with LC3 and focal expression of nitroY co-localizing with UCHL1 in the gastric myenteric ganglions, indicating possible existence of neuronal excretosomes (Supplementary Fig. 15). In addition, exopher-like membranous structures with no nuclear staining were also observed in the vicinity of gastric myenteric ganglions by S100, αSyn and FH immunostaining.

## Discussion

In summary, we showed that HT22 cells use an excretion system composed of excretosomes, nanotubes and exophers to excrete different type of metabolic wastes with MDA, 4HNE, nitrotyrosine modifications or ubiquitin or LC3-tagged molecules, or lysosomal residues both constitutively at basal oxidative respiration condition and also inducibly under elevated oxidative stress conditions. Experiments also showed that U251 cells excreted LC3 or MDA in exophers under oxidative stress conditions similarly. Lipid peroxidation and nitrotyrosine modifications are important risk factors for neurodegeneration^5, 6^, while we have limited knowledge about the detoxification mechanism of lipid peroxidation or nitration products^25, 26^. The current study established a clear link between autophagy with lipid peroxidation and nitration modification removal as excretosomes and exophers contain high concentrations of metabolic wastes such as MDA, 4HNE and nitroY that are enriched in autophagic vesicles. The formation of excretosomes and exophers, which could be biochemically equivalent structures, are linked to a microtubule-directed autophagic vesicle clustering and fusion process with also co-clustering of other related vesicles such as lysosomes and MVBs. LC3 and autophagosome association with microtubules has been previously reported^27, 28^. Metabolic waste degradation and excretion is likely achieved in mobile cellular domains on microtubule tracks and with collective interactions between autophagosomes, MVBs and lysosomes. Histological evidence from the gastric myenteric ganglions supported the existence of neuronal excretosomes and exophers although more experiments especially real-time imaging are needed to characterize neuronal excretosomes and exophers *in vivo*. The proteostatic balance of the cells appears to be maintained not only by synthesis and degradation but also by excretion. H_2_O_2_ oxidative stress induces HT22 cell death with multiple defects including excretion failure, oxidative DNA damage, cytoskeleton disruption, glycolytic enzyme and stress response deficiency but not Aβ, αSyn, phos-TAU accumulation. Caspase-3, nuclear p38 and p53 activation could be also linked with H_2_O_2_-induced HT22 cell death. Activated Caspase-3 and phos-p38 also existed in exophers, which might explain why many extracellularly deposited protein aggregations are fragmented by proteinases or modified with phosphorylation. Previously, autophagy, lysosome and ubiquitin had been shown all linked with secretion^29-31^. In light of recent research progress, many of these “secretion” events could be preceded by an unconventional “excretion” step by excretosomes and exophers. The enclosed small vesicles such as exosomes or cargo proteins could be released from excretosomes and exophers through secondary events as excretosomes and exophers possess some biological activities by themselves even without the instructions from the nuclei.

Intracellular autophagic vesicle clustering was previously described in a published study related to neuronal Herpes Simplex Virus infection, however in this paper, the authors stated autophagic vesicle clustering only happened in virus-infected cells but not in normal cells^32^. Our study indicated that there is a constitutive microtubule-directed autophagic vesicle clustering process to degrade and to excrete metabolic waste materials in wild type HT22 cells. The extracellular disposal of membrane enclosed MVB-like small vesicle clusters has recently described by a number of studies^33, 34^. One of these studies with colorectal carcinoma cells provided evidence that the clustered small vesicle release relates to LC3 and RAB7 signaling^34^. Cell migration and nanotube-like retraction fiber breakage was proposed as a releasing mechanism for migrasomes, a type of multivesicular extracellular vesicles^33^, which might help to explain the phenomenon of excretosome release at cellular nanotube terminals. Since both nanotube terminal membrane breakage and membrane blebbings might be related to calcium signaling, we propose that calcium could be a regulator of excretosome or exopher disposal. The positive phos-CAMK2 staining in excretosomes supported this hypothesis. This study indicated that multivesicular cargo release likely represent the events of the disposal of metabolic wastes through autophagic vesicle clustering. To our knowledge, we also reported for the first time, the extracellular disposal of large specialized autophagic vacuoles with simultaneous waste degradation and excretion functions in mammalian cells. It is well-known that large autophagic vacuoles exist in the yeast cells to perform degradative functions^35^. Although the yeast vacuole is a constitutive component of yeast diploid life cycle, however, these vacuoles can be excreted during yeast meiosis and spore formation^36^. Thus, an excretory function of large autophagic vacuoles might be evolutionarily conserved.

It is not clear about the roles of Aβ, TAU, αSyn in autophagic vesicle clustering and fusion. However, αSyn was shown to be important for synaptic vesicle clustering^37^. αSyn overexpression induced cytoplasmic vesicle aggregations in human U251 cells^38^. The participation of αSyn in autophagic vesicle clustering regulation is very likely. Aβ has also been shown to preserve membrane fusion properties that could be important for autophagic vesicle clustering and fusion^39^. We detected TAU in the enlarged autophagic vacuole. However, there was little past literature indicating TAU’s role in vesicle clustering or fusion except that TAU was reported to be existing in exosomes^40^. On the one hand, the aggregates of Aβ, TAU, αSyn could be target cargos that are removed through autophagic process. On the other hand, the aggregation properties of Aβ, TAU, αSyn might facilitate autophagic vesicle clustering, fusion and metabolic waste removal process. Previous studies indicated that Aβ, αSyn and TAU have important physiological functions^41-43^ and cell survival could be affected adversely either in their absence or when they are being massively produced. The basic functions in normal neuron cells of Aβ, αSyn, TAU need to be vigorously examined before we can truly clarify their roles, diagnostic and therapeutic potentials in Alzheimer’s and Parkinson disease^44^.

Excretosomes and exophers likely to have unstable membrane structures since they contain highly concentrated metabolic wastes and they will readily release their contents after membrane breakages. In view of previous literature, there is a long history to study membrane blebbings and exopher-like structures previously named as ectosomes^45^, microparticles^46^, cell fragments^47^, cytoplasts^48^ or microplasts^49^ in different cell types and under different experiment conditions, indicating a general mechanism for cells responding to stress, which is not limited to neuronal cells. We think excretosomes and exophers are still proper names to use because they are assigned with an important and clear waste excretion function. This study also showed that the drastic reduction of glycolytic proteins such as GAPDH, PKM2, ENO1 in dying H_2_O_2_-treated cells, suggesting glycolysis defects likely contribute to neurodegeneration upon oxidative stress, which is consistent with previous reports indicating lower aerobic glycolysis activity in Alzheimer’s disease models^50, 51^. As many different kinds of metabolic products are released at high quantities into exophers by stressed cells, exopher-genesis likely have a great impact on amyloidogenic and prion diseases, sterile inflammation, aging associated secretory phenotype, which are all important topics of neurodegenerative disease studies. Furthermore, the high amount of proteins released from exophers could also become self-antigens if not cleaned up quickly from the system. We believed that exopher-released autoantigens can explain human autoimmune diseases in many occasions, which could not be sufficiently addressed by cross-reactivities between self-proteins and pathogen antigens. The Calcein AM dye transmission experiments suggest that neuronal stress signals might spread to neighboring neurons or glial cells through nanotube and exopher networks, which could partially explain the progressively worsening phenotypes of many neurodegenerative diseases, such as Alzheimer’s disease, Parkinson’s disease, ALS and prion diseases. Based on the experiments in this paper and previous research, we draw a hypothetic scheme to show the basic framework of a cellular excretion system in Fig. 8g. The study of the cell excretion regulation likely will stimulate fresh thinking on the causes, diagnosis and treatments of many of these complicated human diseases.

## Materials and Methods

### List of antibodies for immunocytochemistry

The following primary antibodies and dilutions have been used in this study: MDA (Abcam ab6463, 1:200), 4HNE (Abcam ab46545, 1:200), nitroY (Sigma N0409, 1:200), UBB (Proteintech 10201-2-AP, 1:200), LAMP2 (Proteintech 66301-1-Ig, 1:100), LC3 (This antibody detects both LC3A and LC3B.) (Proteintech 66139-1-Ig, 1:100), LC3B (This antibody detects only LC3B) (CST 3868,1:200), CD9(Proteintech 60232-1-Ig, 1:200), CD63(Abcam ab134045, 1:200), GAPDH(Proteintech 10494-1-AP, 1:200), beta Actin(Proteintech 20536-1-AP, 1:200), alpha-Tubulin(Proteintech 11224-1-AP, 1:200), phos-CAMK2 (detecting phos-CAMK2α and phos-CAMK2β) (Abcam ab124880, 1:200), phos-TAU (Abcam ab151559, 1:200), G3BP1 (Abcam ab181150, 1:200), Hsp70 (Abcam ab181606, 1:200), AnnexinA2 (Abcam ab178677, 1:200), PKM2(CST 4053, 1:200), ENO1 (Proteintech 55237-1-AP, 1:200), Aβ (Abcam ab201061, 1:200), αSyn (Proteintech 10842-1-AP, 1:200), phos-p53 (CST 12571, 1:200), 8-OHdG (Santa Cruz sc-66036, 1:100). The following secondary antibodies and dilutions have been used in the study: donkey anti-mouse Alexa 594nm secondary antibody(Jackson ImmunoResearch, 715-585-150, 1:400), donkey anti-rabbit Alexa-488nm secondary antibody(Jackson ImmunoResearch, 711-545-152, 1:400), donkey anti-rabbit Alexa 594nm secondary antibody(Jackson ImmunoResearch, 711-585-152, 1:400), donkey anti-mouse Alexa-488nm secondary antibody(Jackson ImmunoResearch, 715-545-150, 1:400).

### Cell culture, H_2_O_2_ treatment and live cell staining

HT22 cell line was obtained as a generous gift from Dr. Qiang Wang of the Department of Anesthesiology, Xijing Hospital, Xi’an, China. Cells are cultured in DMEM high glucose medium (Corning, USA) supplemented with 10% heat-inactivated fetal bovine serum (Tianhang Biotechnology), 100 U/ml penicillin and 100 ug/ml streptomycin (Beyotime Biotechnology). Cells are plated on Poly-L-lysine (Sigma P4832)-coated coverslips in 24-well tissue culture plates at approximately 40% confluency in 0.5 ml culture medium. 1 mM H_2_O_2_ (1 μl from a 500 mM H_2_O_2_ stock for 0.5 ml culture medium) was added after plating for 2hrs, and the incubation continued for 4hrs. In the controls, 1μl H_2_O was added. 1 mM H_2_O_2_ treatment of HT22 cells for 4 hours induce roughly 10% cell death calculated from PI-Hoechst33342 staining. Live cell staining is performed with the following conditions: PI (Sigma P4170, 10 μg/mL for 15 minutes), Hoechst (Sigma B2261, 1 μg/mL for 15 minutes), Calcein AM (Yeasen Biotech., 2.5 μg/mL for 30 minutes), MitoTracker (Molecular probes M7512, 200 nM for 30 minutes). All these dyes are diluted with cell culture medium at the indicated concentration and incubated with cells at 37 degree in the cell culture incubator for the desired time period. When performing autophagy inhibition experiment, Chloroquine phosphate (SantaCruz, sc-205629) was used at 100 μM concentration.

### Dye transmission experiment

HT22 cells were first labeled with Calcein AM for 30 minutes. Then, labeled cells were trypsinized and mixed with trypsinized unlabeled cells at 1:6 dilution and replated. The co-culture continued for 5 hours and then stained with Hoechst and imaged.

**Immunocytochemistry** was performed as described^52^. Briefly, cell culture medium was removed after culturing. The cells on coverslips were rinsed with PBS briefly for 2 times. Then, the cells were fixed in 4% PFA for 20 minutes at room temperature. The cells were rinsed again with PBS twice and proceeded for TBST 3% BSA blocking. After blocking, the samples were incubated with primary antibodies at room temperature for 2 hrs. followed by 5 washes of TBST. After that, the samples were incubated with fluorescent secondary antibodies overnight at 4 degrees. The treated samples were washed again with TBST 5 times the second day and mounted with PBS+50% glycerol supplemented with H33342 nuclear dye and ready for imaging. For 8-OHdG staining, an additional step of 2 M HCl pretreatment of cultured coverslips for 30 minutes at 37°C was performed.

### Imaging and morphometry analysis

All images were taken with a Leica DM6000B microscope (Leica Microsystems, Wetzlar, Germany) with A4, L5 and N3 fluorescent filter cubes. The microscope is equipped with 10X eyepiece, and 5X, 10X, 20X, 40X objective lens and 1.6X digital zoom. For time lapse imaging, OCam screen capturing software was used. The obtained movie files were compressed with Quick Media Editor movie editing software. When comparing marker immunostaining densities, identical exposure settings for fluorescent channels was used for both control and test groups. The exposure time for multiple marker co-localization studies were longer and sometimes overexposed comparing to the density comparison studies in order to show the fine details of the staining. Images were then analyzed with Lecia Advanced Fluorescence software. Draw counter tool was used for cell counting. ROI tools were used to select subjects of interest. Average area density was used as the parameter to define the marker densities of excretosomes, exophers and cell bodies. Approximately 10 to 20 control subjects were first measured with the software and the average value was used as the control value. Around 10 to 20 experiment subjects were measured with software and the obtained values divide the control value to obtain the values of fold of changes. The fold of changes was then expressed as means with standard deviations. The density analysis was carried out under conditions that the categories of the samples were blinded to the observers. For the size measurement of excretosomes, the length measurement tool of Image J software was used with a microscope scale. Quantitation in the figures showed one set of experiments only. All experiments have been repeated multiple times to ensure that the conclusions were reproducible.

### Statistics

All data first went through a Shapiro & Wilk normality test using SPSS Statistics 19 software. Two-tailed unpaired T-test was used to compare the means of data with a normal distribution with Excel 2007 software. For data that were not normally distributed, nonparametric Mann/Whitney test was performed to compare the means with Megastat addin version 10.2. The p Value threshold for statistical significance is set at 0.05. If p<0.05, then the difference was considered statistically significant. when p<0.001, it was labeled as p<0.001. If p≧0.001, then the p Value was shown as it was. The degree of freedom (df) was always shown. “*” in the statistical figures indicated the differences are statistically significant.

## Data and materials availability

All data needed to evaluate the conclusions in this paper are present either in the main text or the extended materials.

## Acknowledgements

We thank Dr. Qiang Wang (Department of Anesthesiology, Xijing Hospital, Xi’an, China) for HT22 cells. We also thank Housheng Wang, Lu Xu, Wenting Xuan, Pingxin Liu, Xiaoyi Bao for excellent lab assistance.

## Author contributions

H.F. conceived the study, H. F. designed and supervised the experiments; H.F. and J. L. performed the experiments; H. F., J. L., D.P., W. J., D. C. did the data analysis; H. F. wrote the manuscript; All authors reviewed the manuscript.

## Supplementary Information

### Supplementary Materials and Methods

#### Antibodies and other reagents

The following additional antibodies have been used in supplementary experiments: Cleaved Caspase-3 (CST 9664, 1:200), phos-p38 (Abcam ab178867, 1:200), Cytochrome C (Abcam ab133504, 1:200), HMGb1 (Abcam ab79823, 1:200), p53 (Proteintech 10442-1-AP, 1:200), S100 (Abcam ab183979 1:500), TAU (Abcam ab32057, 1:500), UCHL1 (Proteintech 66230-1-Ig, 1:500). The tissue slides used in immunohistochemistry experiments for research purpose were obtained from Shanghai ENOVO Industrial Co.

#### Immunohistochemistry

Paraffin sections of human gastritis tissues were deparaffinized with toluene and rehydrated with an inverse ethanol gradient following standard procedures. Sections were then treated with 10 mM pH 6.0 sodium citrate antigen retrieval solutions at boiling condition for 20 minutes with microwave heating. After cooling down, the sections were then stained with antibodies with the same procedure as for immunocytochemistry of cultured cells.

#### Cell culture

U251 cells were purchased from the Chinese Academy of Sciences Cell Bank. U251 cells were treated with 2 mM H_2_O_2_ for 6hrs to induce cell death or at a mild stress condition with 1 mM H_2_O_2_ for 3hrs without inducing cell death. Other experiment conditions were the same as for the HT22 cell experiments.

#### Time lapse imaging file editing

The time-lapse imaging file recorded from the HT22 hydrogen peroxide treatment experiments were clipped and compressed 8X to show the exopher formation process starting from membrane blebbing by using the Quick Editing movie editing software.

---

**Supplementary Fig. 1.**
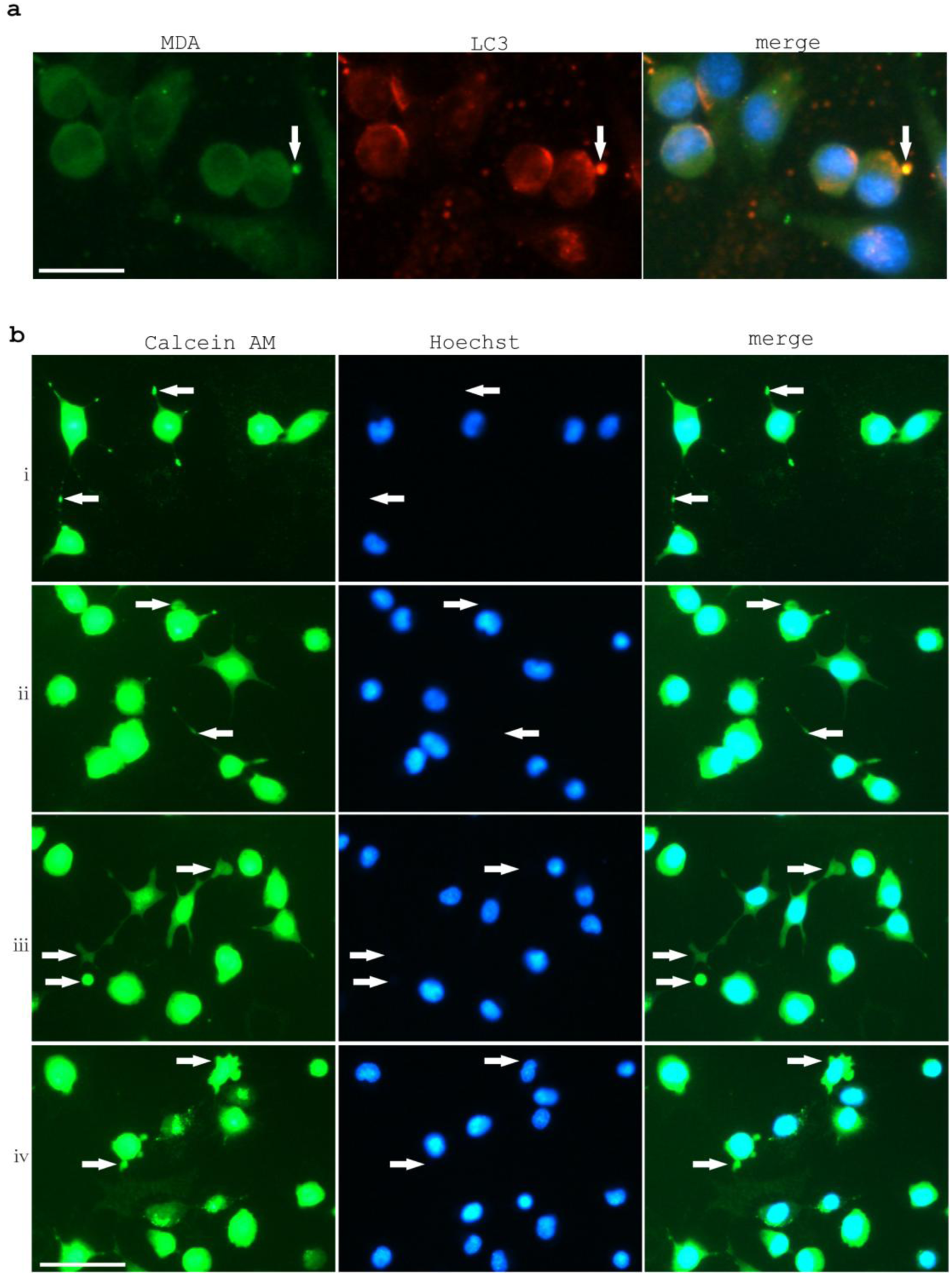
Different forms of excretosomes, nanotubes and exophers in HT22 cells. **a.** Sometimes excretosomes seem to derive from small plasma membrane blebbings in control untreated HT22 cells. The arrow indicated a membrane blebbing form of excretosome positive with both MDA and LC3 immunostaining. Scale bar, 25 μm. **b.** Calcein AM staining of excretosomes, nanotube, and exophers in H_2_O_2_-treated HT22 cells. (i), Calcein AM labeling in nanotubes and excretosomes. (ii), Calcein AM labeling in membrane blebbings and excretosomes. (iii), Calcein AM labeling in exophers. (iv), Calcein AM labeling in membrane blebbings. Arrows indicate excretosomes, nanotubes and exophers. Scale bar, 50 μm.

**Supplementary Fig. 2.**
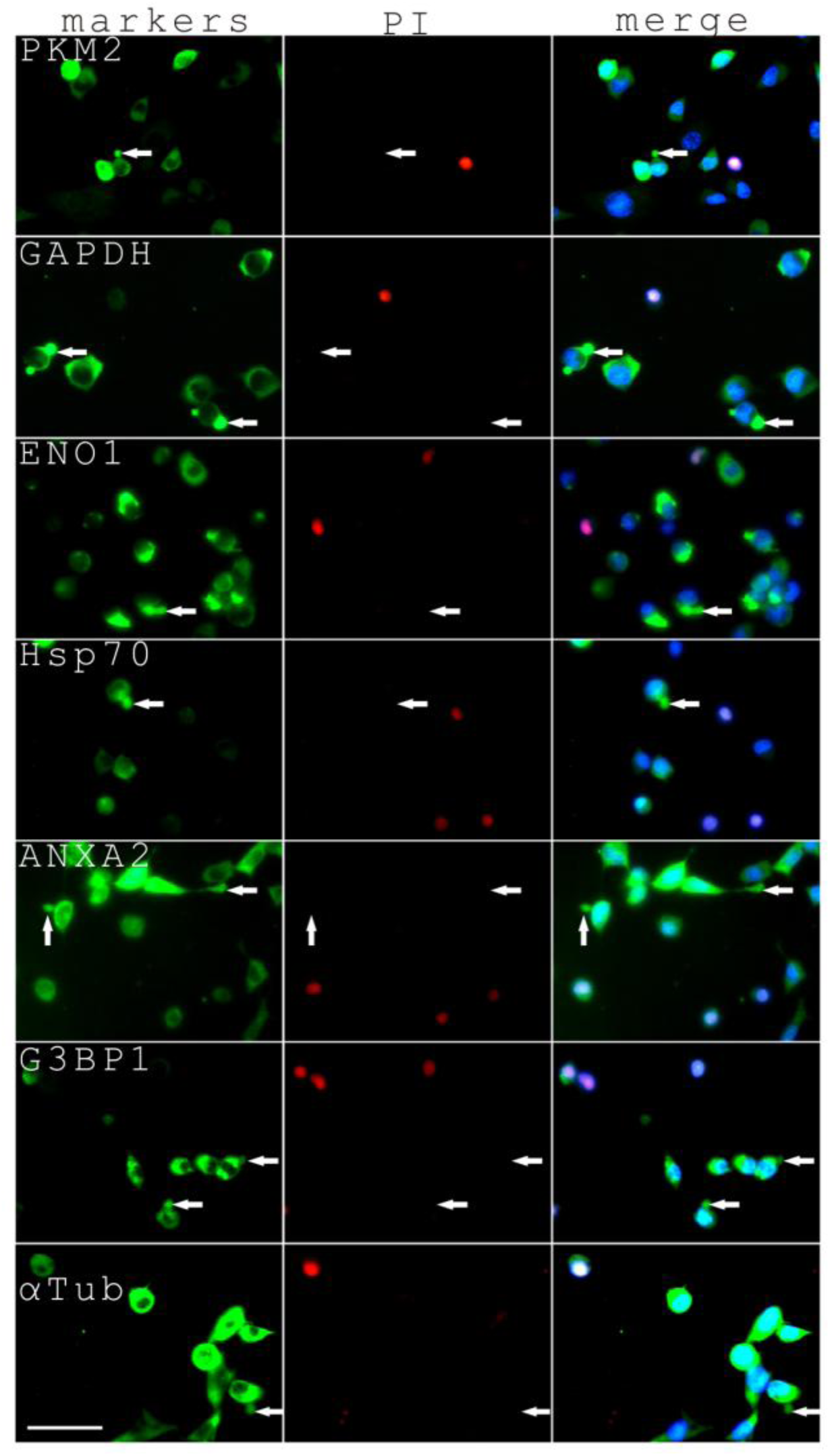
Excretion of PKM2, GAPDH, ENO1, Hsp70, ANXA2, G3BP1, αTub in exophers by live H_2_O_2_-stressed HT22 cells. Arrows indicated exopher formation from PI-negative cells. Scale bar, 50 μm.

**Supplementary Fig. 3.**
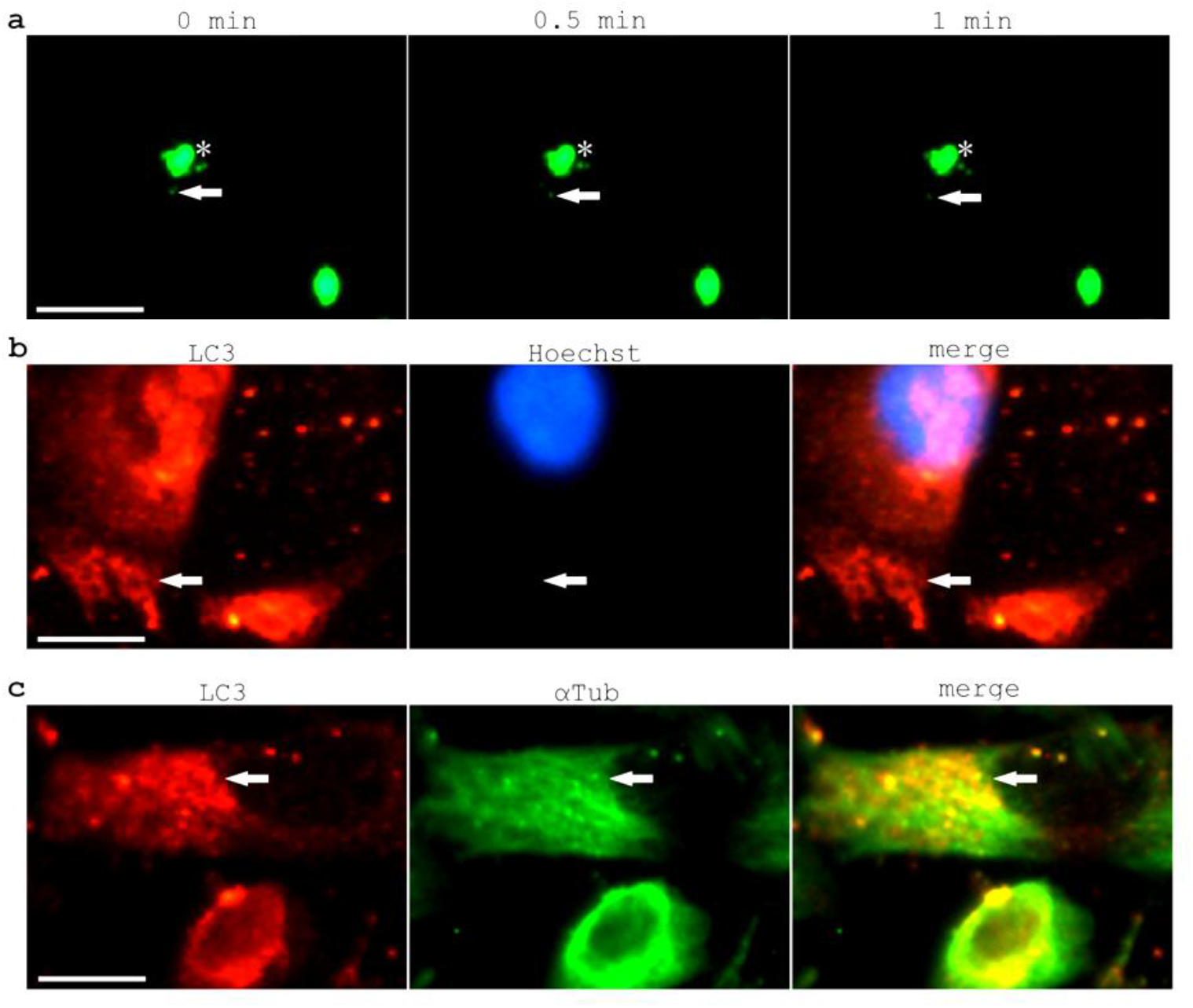
Autophagic vesicle clustering associates with microtubules. **a.** A real-time observation on vesicle movement by Calcein AM labeling in a one-minute time frame. Scale bar, 50 μm. **b.** A magnified image of autophagic vesicle clustering at the cell periphery by LC3 immunostaining. Scale bar, 12.5 μm. **c.** A magnified image of extensive co-localization of the microtubule protein αTub and autophagic vesicle staining. Scale bar, 12.5 μm

**Supplementary Fig. 4.**
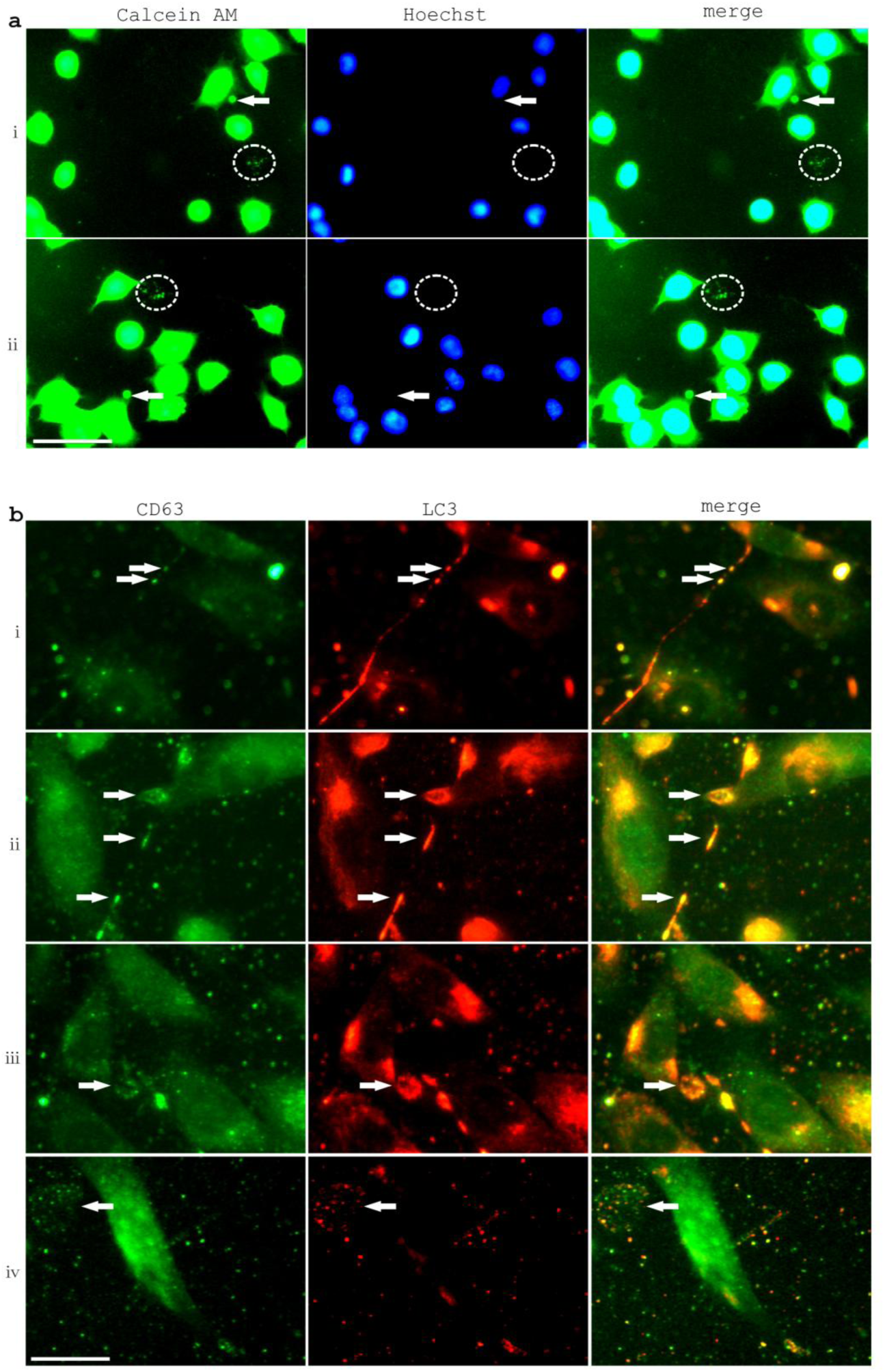
Multivesicular properties of excretosomes in wild type HT22 cells. **a.** Calcein AM-labeled excretosomes with compacted signals (marked with asterisks) and with multivesicular forms (marked with circles). Scale bar, 50 μm. **b.** Multivesicular features of excretosomes were shown by CD63 and LC3 immunostaining with magnified pictures. The pictures emphasized the frequent but not complete CD63-LC3 co-localization in autophagic vesicle clustering in nanotubes (i), in an array of autophagic vesicle clusters (ii), in a large excreted autophagic vacuole (iii) and in a large less-condensed excretosome (iv). Scale bar, 25 μm.

**Supplementary Fig. 5.**
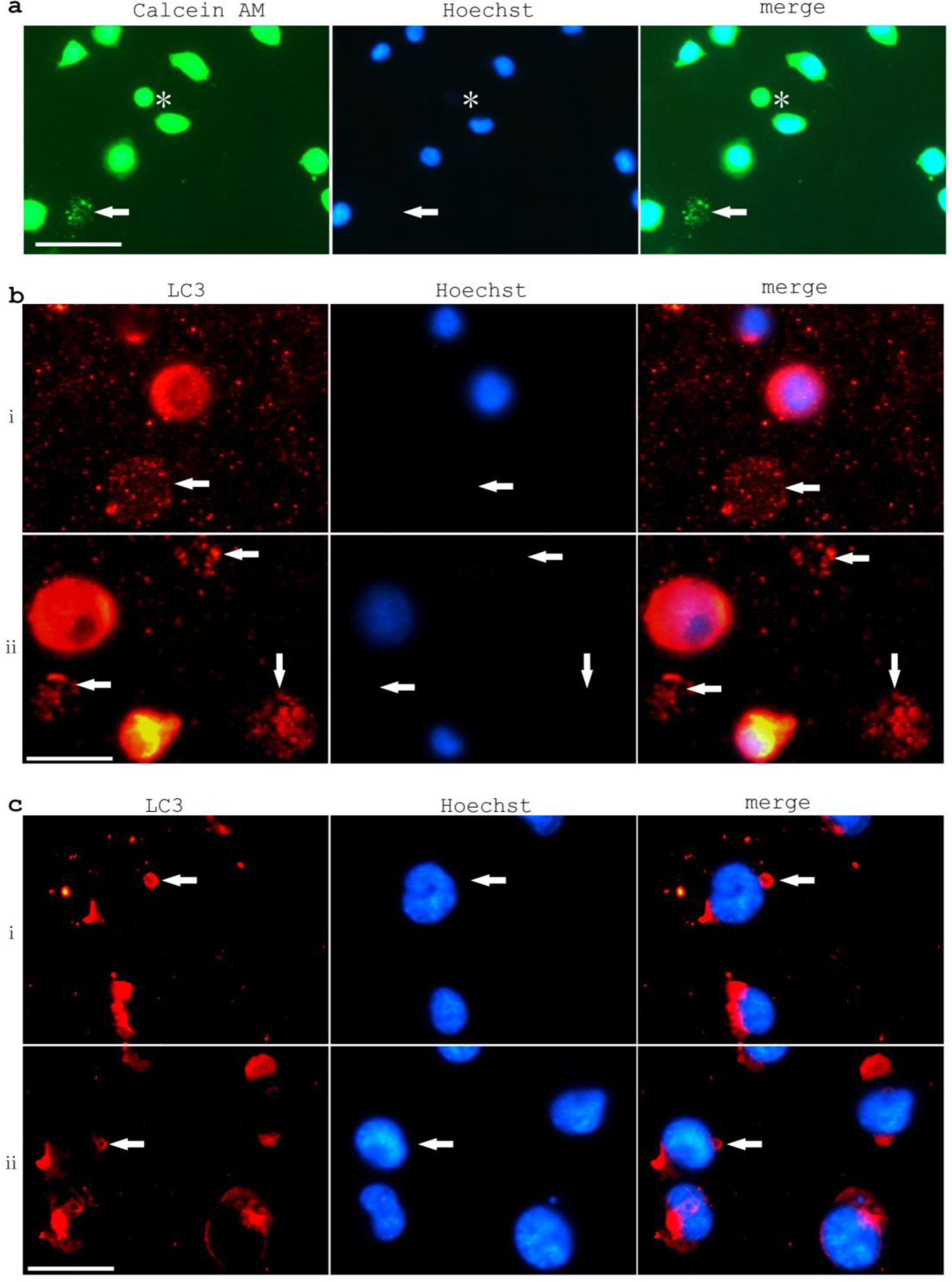
Multivesicular properties of exophers in H_2_O_2_-treated HT22 cells. **a.** Calcein AM-labeled exophers with condensedly compacted signals (marked with asterisk) and with multivesicular forms (marked with arrow). Scale bar, 50 μm. **b.** Multivesicular features of exophers were shown by LC3 immunostaining (marked with arrows). Scale bar, 25 μm. **c.** Fused enlarged autophagic vacuoles indicated by LC3 immunostaining could be observed in some of membrane blebbings during H_2_O_2_ treatment (marked with arrows). Scale bar, 25 μm.

**Supplementary Fig. 6.**
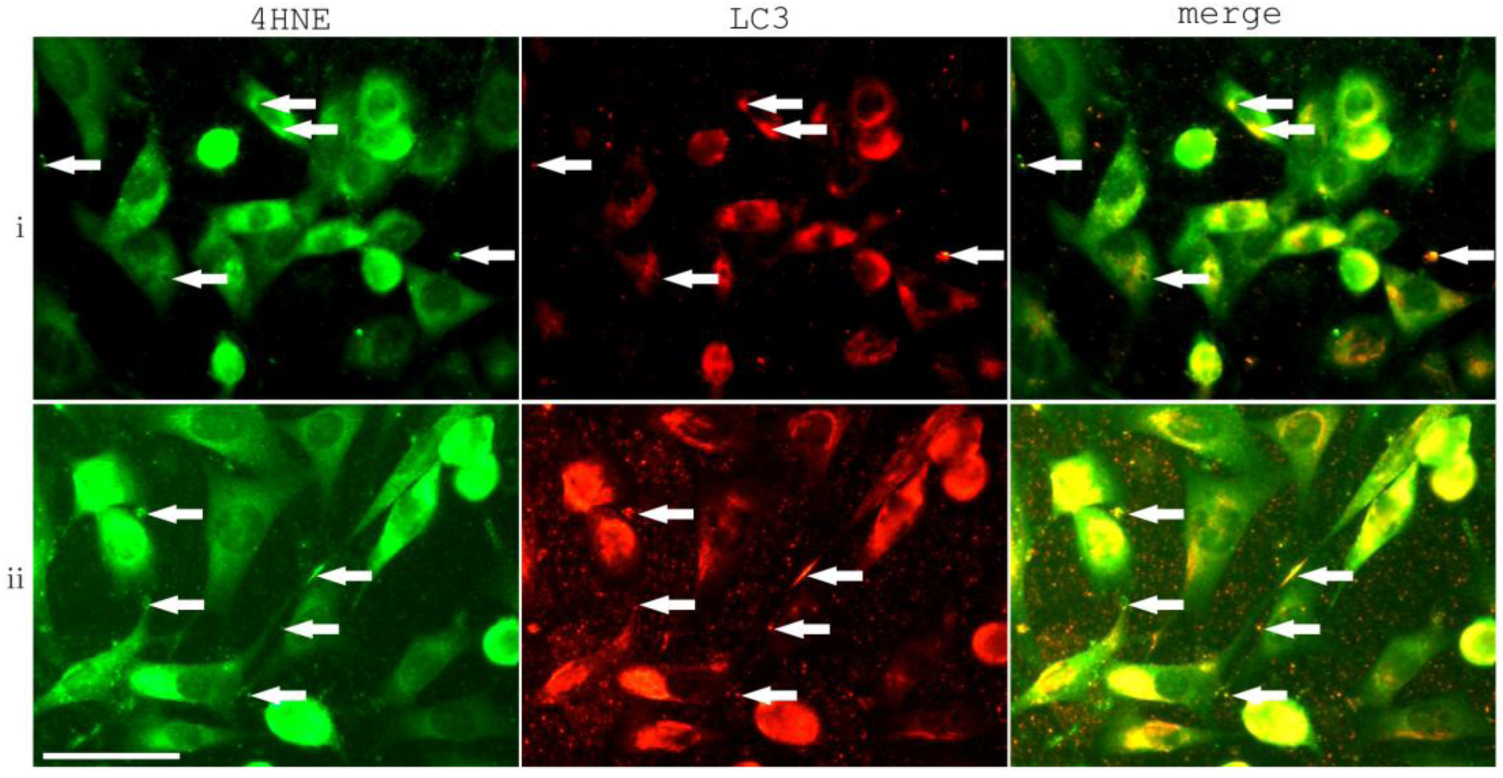
4HNE signals co-localized with autophagic vesicle clustering and fusion in HT22 cells. **i.** 4HNE and LC3 co-labeling in excretosomes and at intracellular vesicle clustering sites. **ii.** NitroY and LC3 co-labeling in excretosomes, and in vesicle clustering in the nanotube and in the enlarged autophagic vacuole. Scale bar, 50 μm.

**Supplementary Fig. 7.**
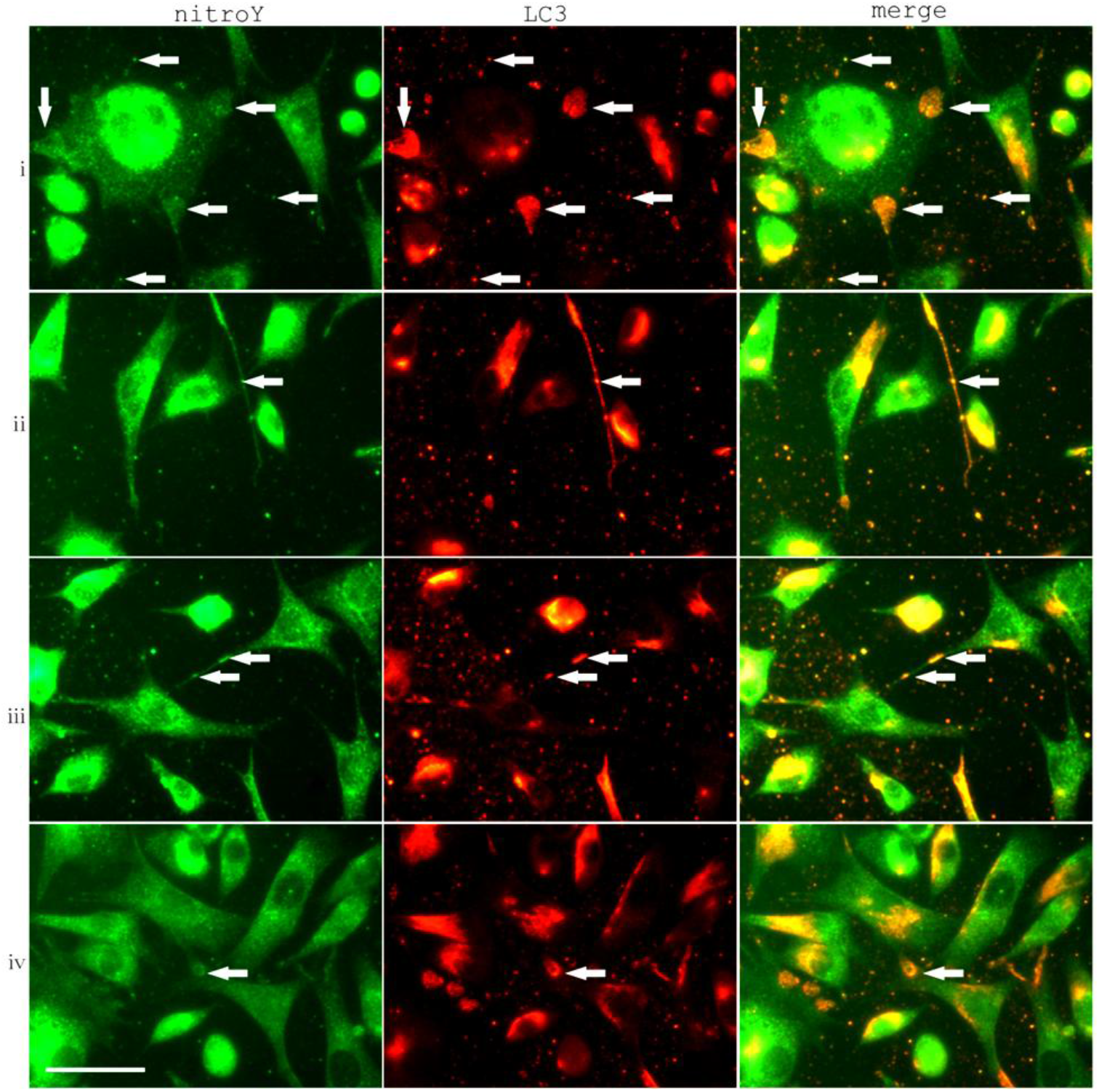
nitroY signals co-localized with autophagic vesicle clustering and fusion in HT22 cells. **i.** nitroY and LC3 co-labeling in excretosomes and at intracellular vesicle clustering sites. **ii.** nitroY and LC3 co-labeling in vesicle clustering in the nanotube. **iii.** nitroY and LC3 co-labeling in an array of autophagic vesicle clusters in the nanotube. **iv.** nitroY and LC3 co-labeling in the enlarged fused autophagic vacuole. Scale bar, 50 μm.

**Supplementary Fig. 8.**
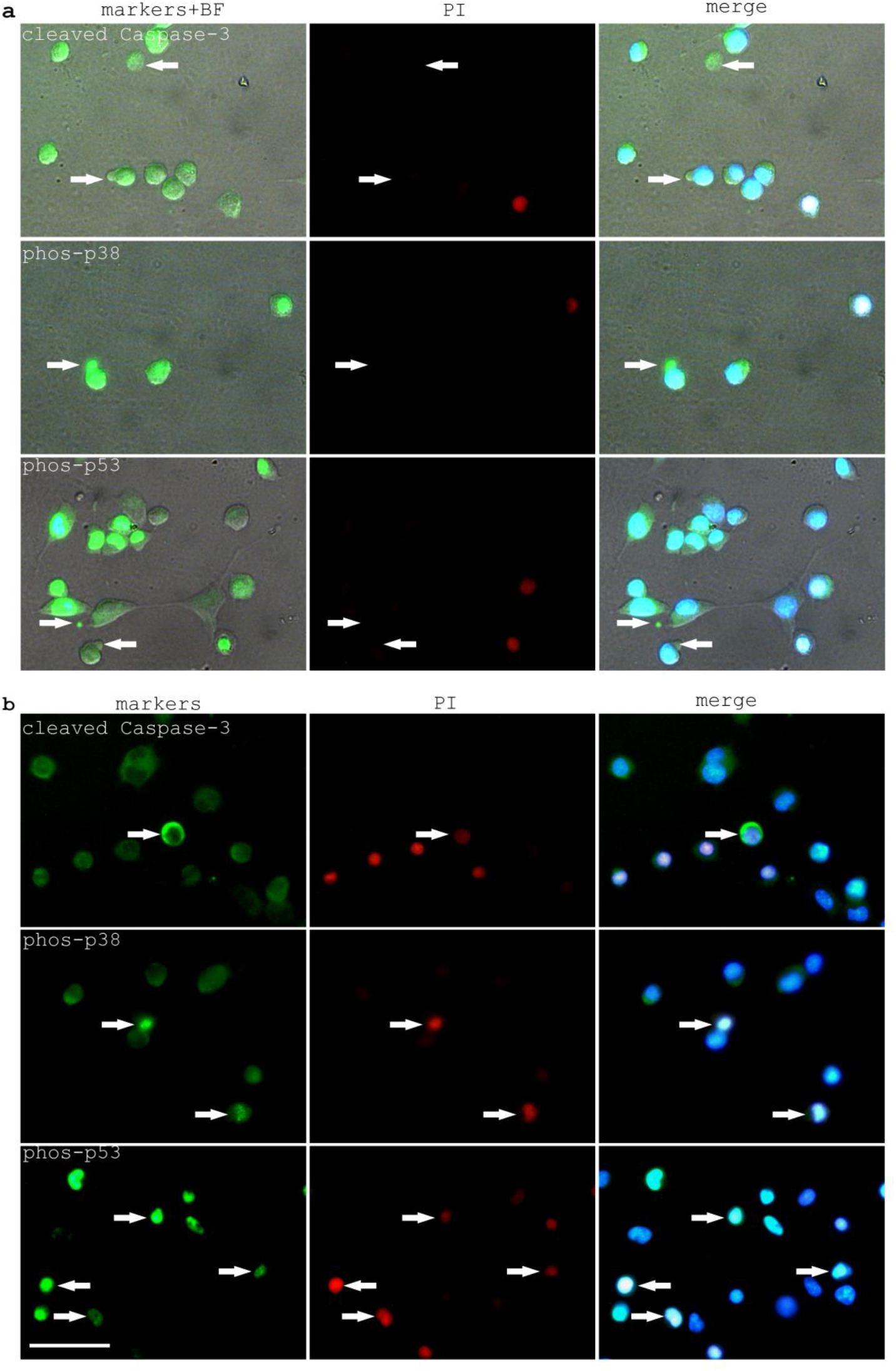
Expression of cleaved Caspase-3, phos-p38 and phos-p53 in H_2_O_2_ treated HT22 cells. **a.** Expression of cleaved Caspase-3, phos-p38 and phos-p53 in exophers of H_2_O_2_ treated HT22 cells. Arrows indicated exopher formation in PI-negative cells. **b.** Cleaved Caspase-3 activation in PI-positive dying HT22 cells and nuclear localization of phos-p38 and phos-p53 in in PI-positive dying HT22 cells. Arrows indicate marker activations in PI-positive cells. Scale bar, 50μm.

**Supplementary Fig. 9.**
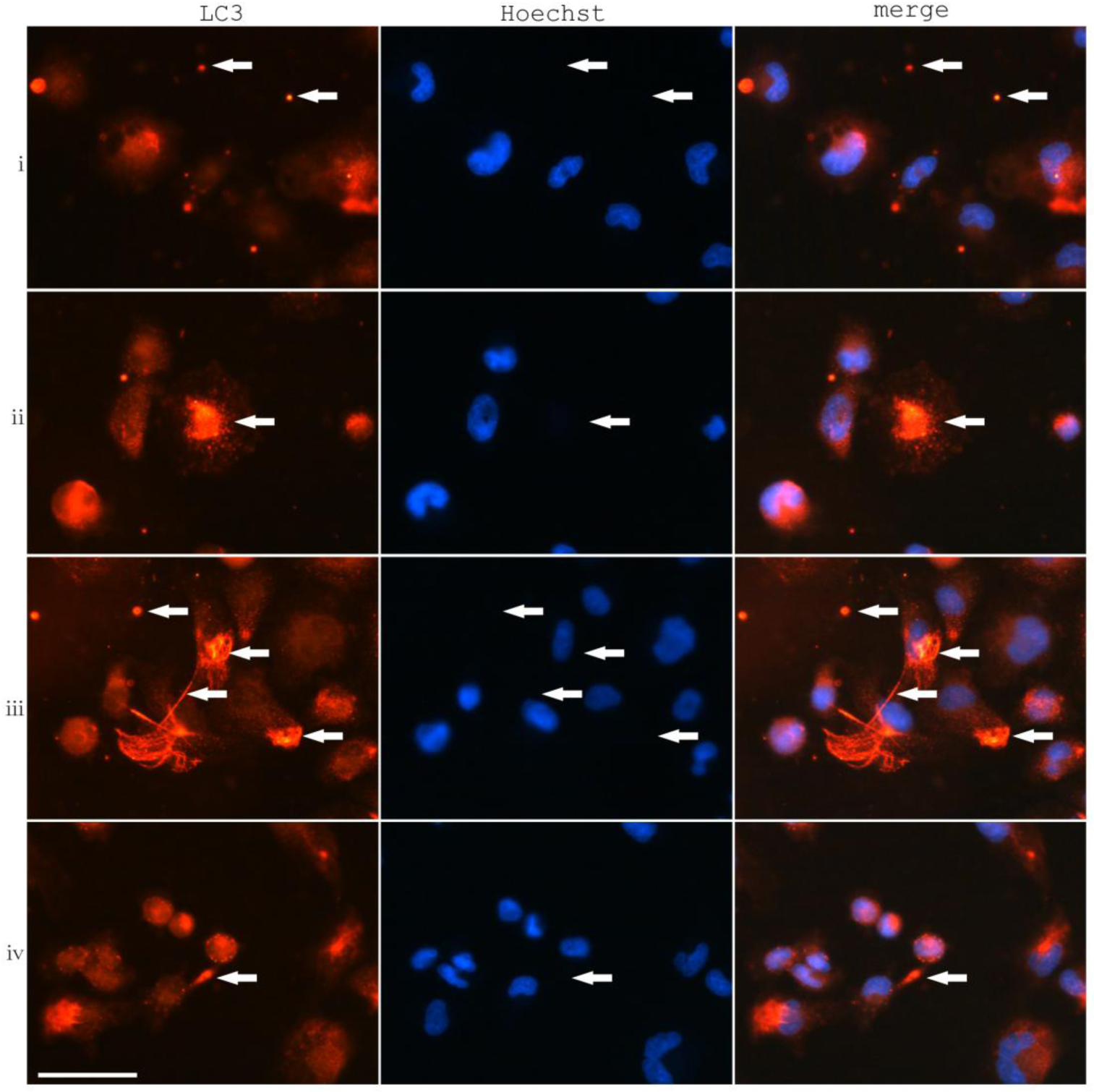
Autophagic vesicle condensations in U251 cells. **i.** LC3 labeling in excretosomes. **ii.** LC3 labeling in a large excretosome. **iii.** LC3 labeling in excretosomes and in intracellular autophagic vesicle clustering possibly associated with microtubule. **iv.** LC3 labeling in autophagic vesicle clustering in the nanotube. Scale bar, 50μm.

**Supplementary Fig. 10.**
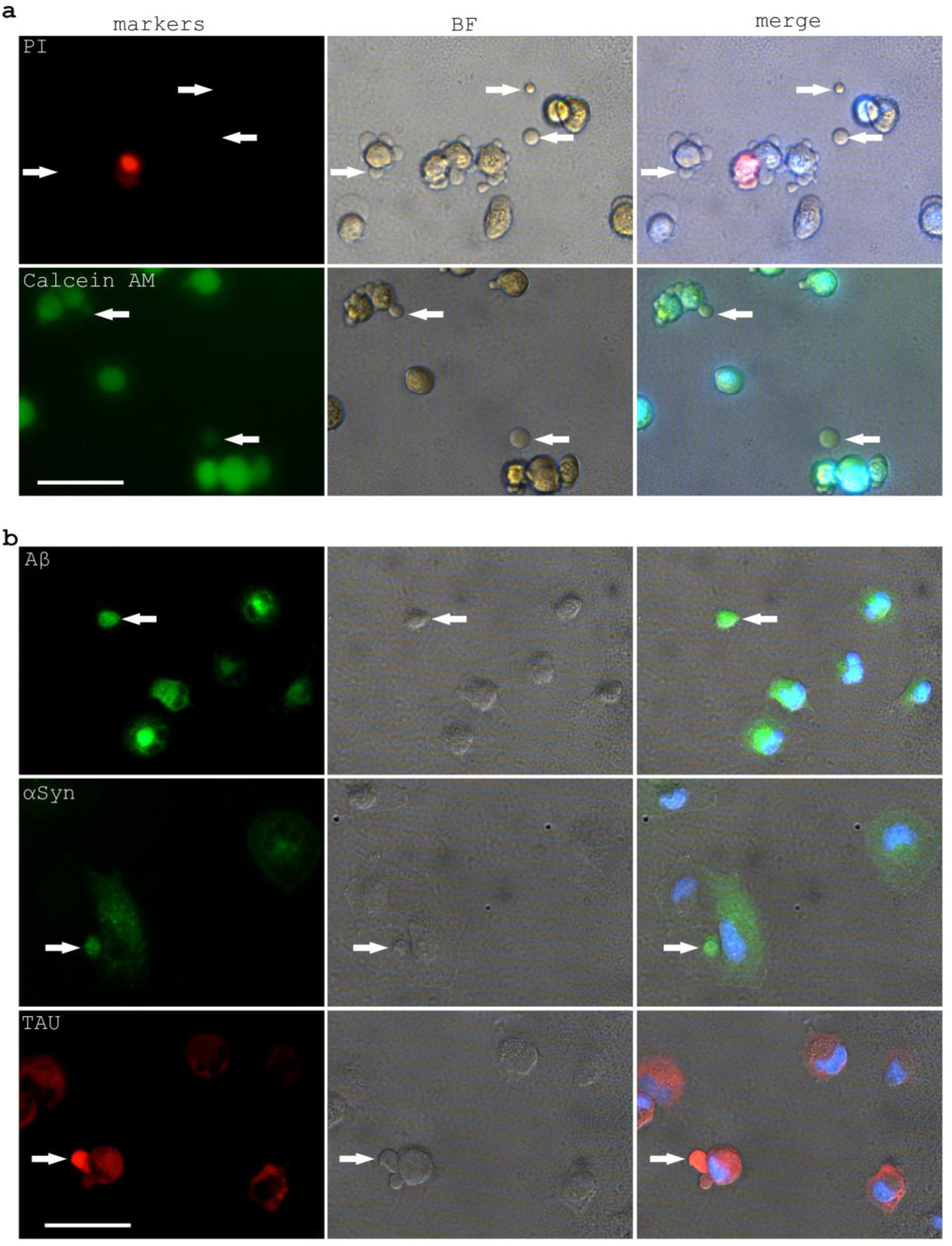
Exopher formation in H_2_O_2_ (2mM 6hrs) treated U251 cells and exopher expression of Aβ, αSyn, TAU in H_2_O_2_ treated U251 cells (1mM 3hrs). **a.** Exopher formation from PI-negative and Calcein AM-positive live U251 cells with H_2_O_2_ (2mM 6hrs) treatment. **b.** Exopher expression of Aβ, αSyn, TAU in H_2_O_2_-treated U251 cells (1mM 3hrs). Arrows point to exophers. Scale bar, 50μm.

**Supplementary Fig. 11.**
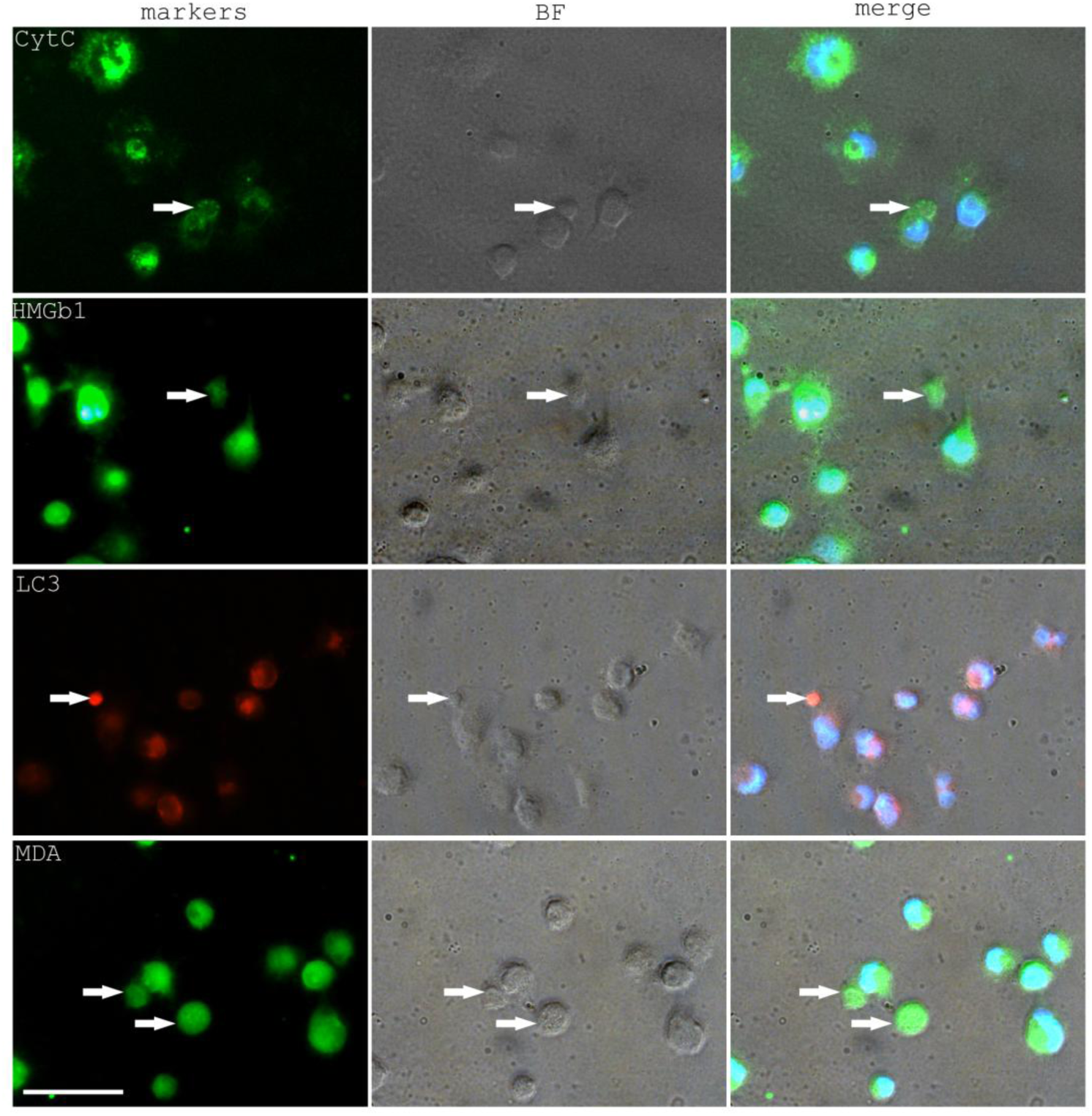
Exopher expression of Cytochrome C (CytC), HMGb1, LC3 in H_2_O_2_ treated U251 cells (1mM 3hrs) and MDA in 2mM H_2_O_2_ treated U251 cells (6hrs). Arrows point to exophers. Scale bar, 50μm.

**Supplementary Fig. 12.**
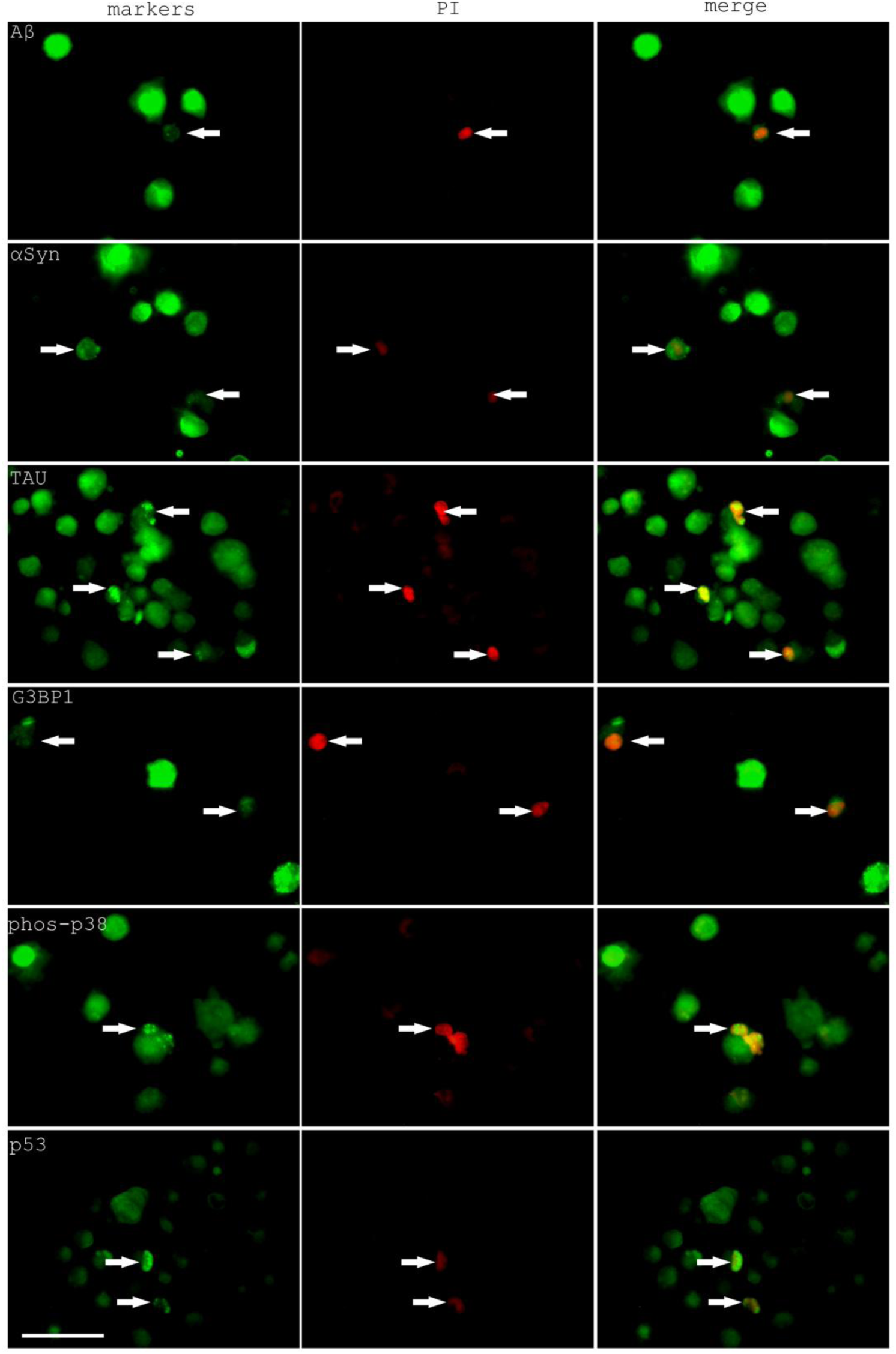
Expression of Aβ, αSyn, TAU, G3BP1, phos-p38, p53 in PI-positive H_2_O_2_ treated U251 cells (2mM 6hrs). Arrows point to marker expression in PI-positive cells. Scale bar, 50μm.

**Supplementary Fig. 13.**
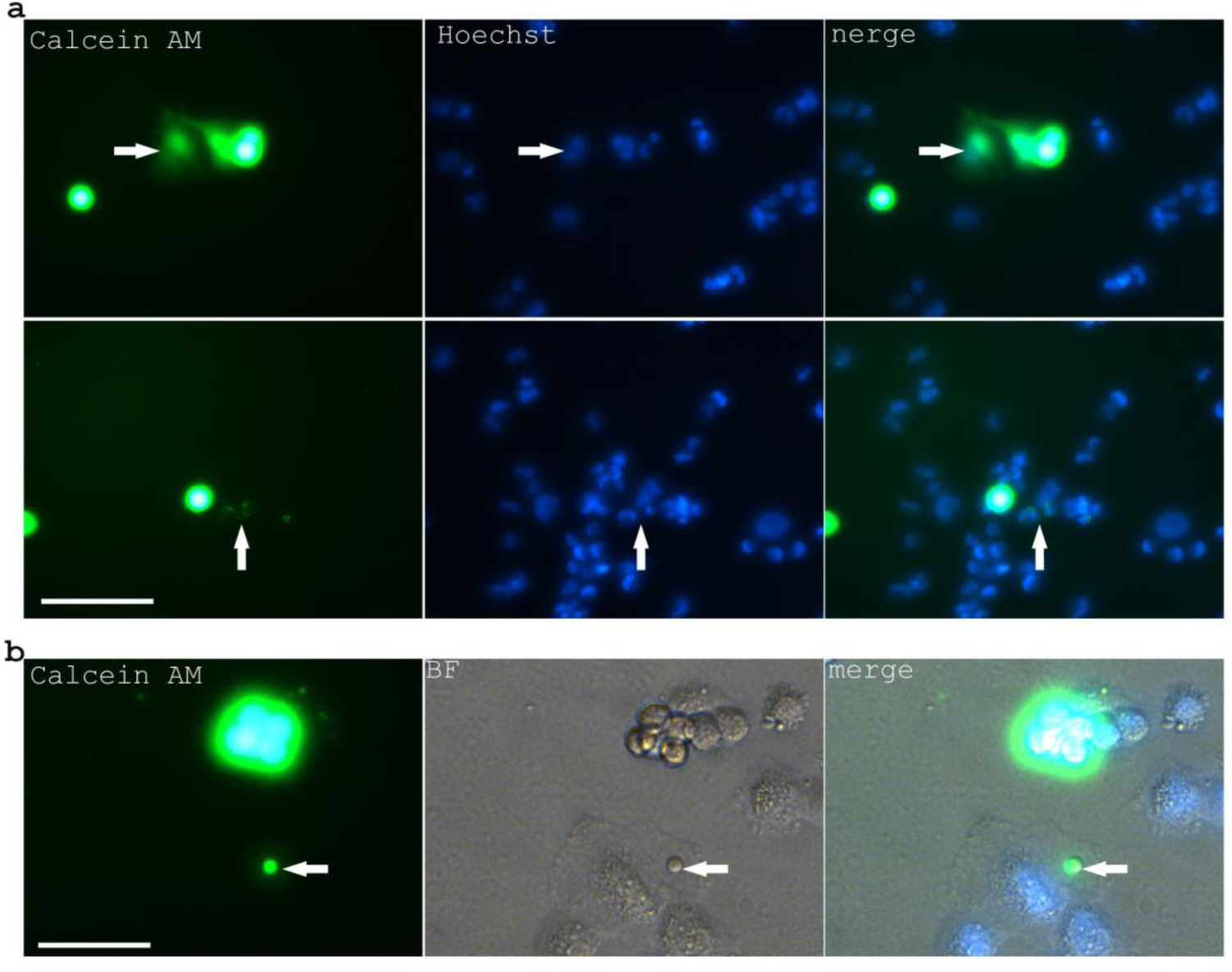
Dye transmission between U251 cells. **a.** Possible dye transmission through cellular nanotubes with diffusive cytoplasmic staining or exophers with granule-like multivesicular staining. Scale bar, 50μm. Arrows indicated recipient cell fluorescence. **b.** Enlarged picture shows the dye transmission through an exopher from a donor cell to a recipient cell. Arrow indicates an exopher. Scale bar, 25 μm.

**Supplementary Fig. 14.**
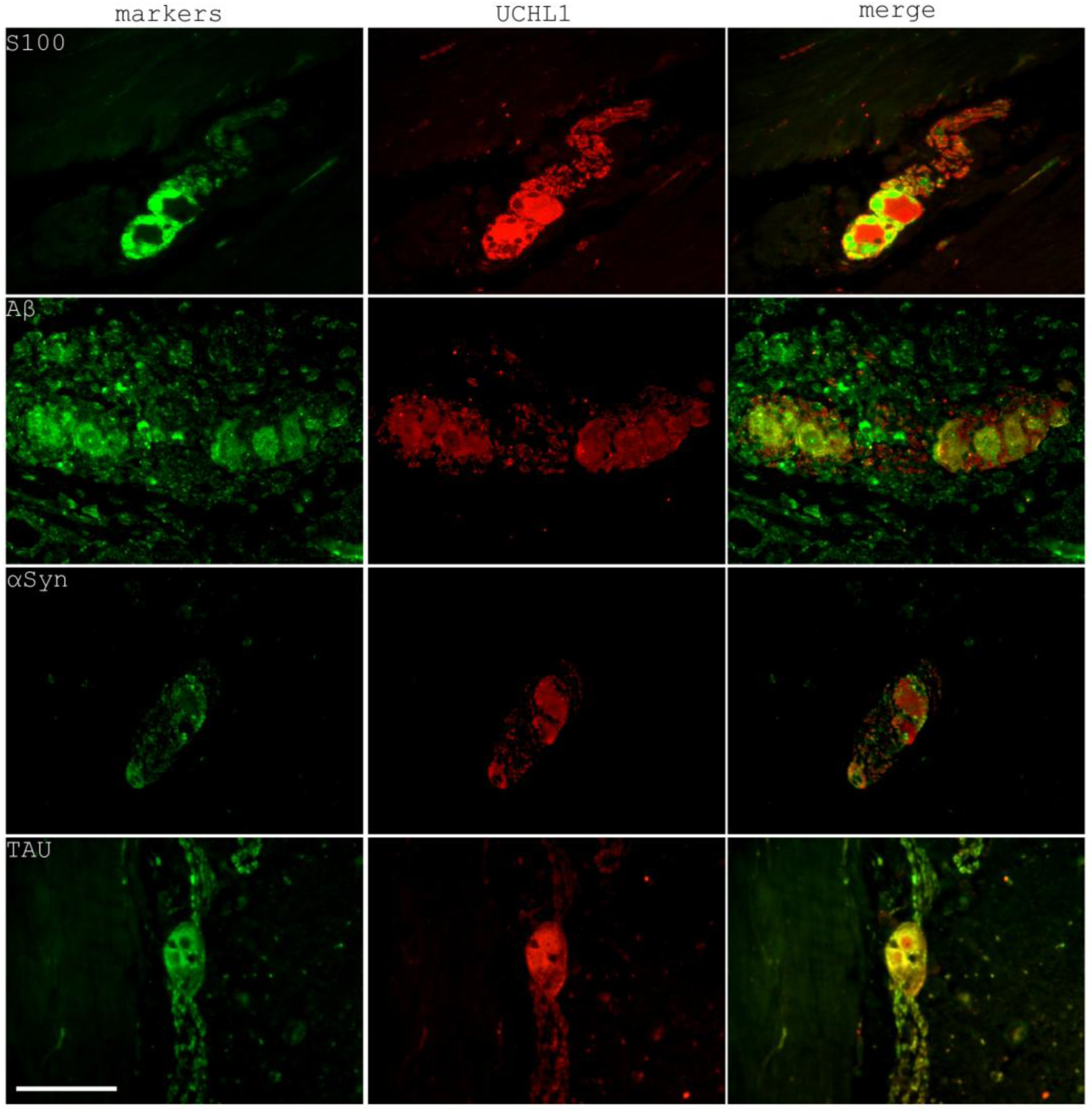
Expression of S100, Aβ, αSyn, TAU in the gastric myenteric plexus with UCHL1 co-staining. Scale bar, 100 μm.

**Supplementary Fig. 15.**
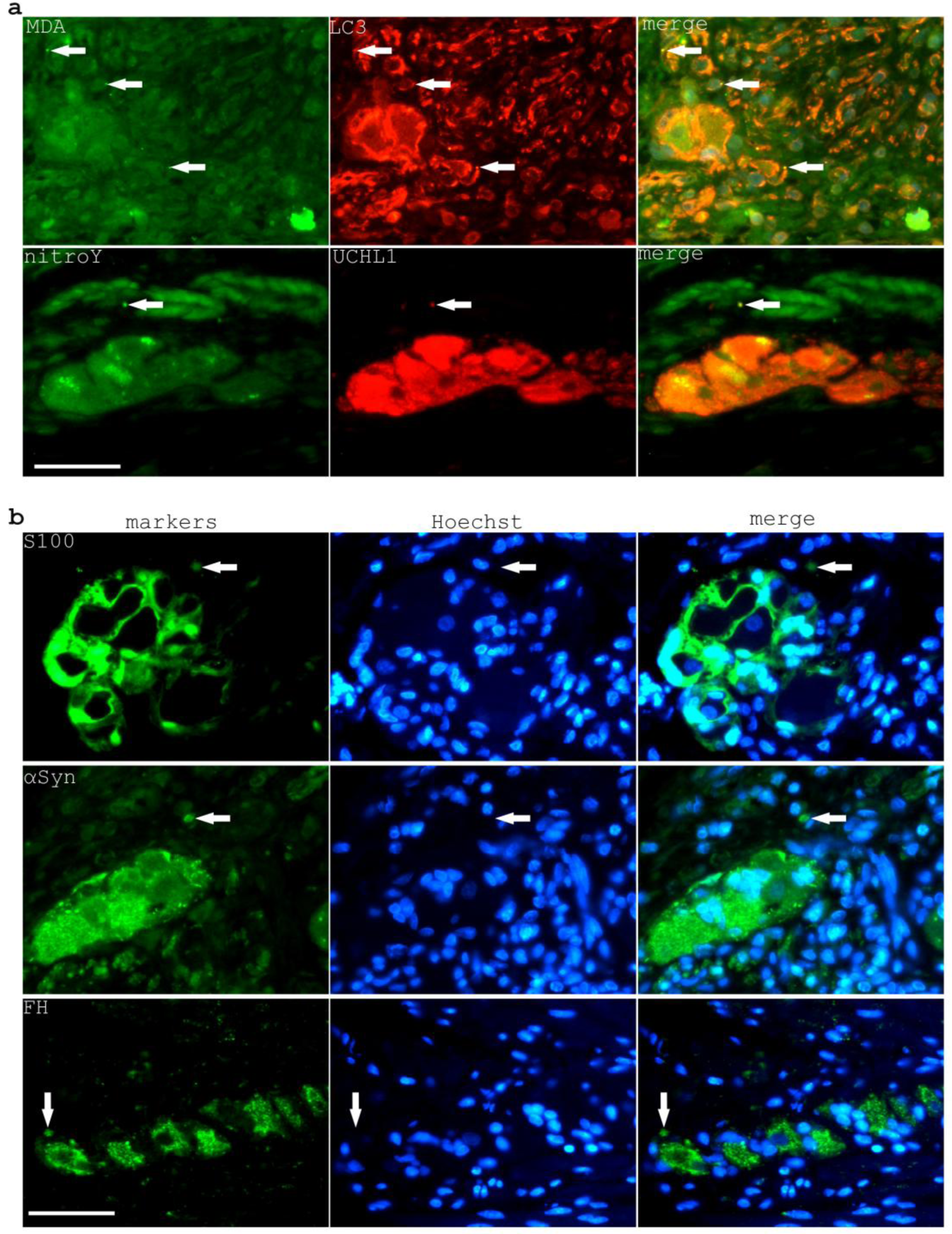
Histological evidence of excretosomes and exophers in the gastric myenteric plexus. **a.** Focal expression of MDA and nitrotyrosine in gastric neurons possibly indicates condensed signals in excretosomes. Arrows indicate focal expressions. **b.** Expression of S100, αSyn and FH in exopher-like structures at the vicinity of gastric neuron ganglions. Arrows indicate possible exopher structures. Scale bar, 50μm.

## References

1 Prince, M. et al. The global prevalence of dementia: a systematic review and metaanalysis. Alzheimer’s & dementia : the journal of the Alzheimer’s Association 9, 63–75 e62, doi:10.1016/j.jalz.2012.11.007 (2013).

2 Chan, K. Y. et al. Epidemiology of Alzheimer’s disease and other forms of dementia in China, 1990-2010: a systematic review and analysis. Lancet 381, 2016–2023, doi:10.1016/S0140-6736(13)60221-4 (2013).

3 Hardy, J. A. & Higgins, G. A. Alzheimer’s disease: the amyloid cascade hypothesis. Science 256, 184–185 (1992).

4 Yan, S. D. et al. Glycated tau protein in Alzheimer disease: a mechanism for induction of oxidant stress. Proceedings of the National Academy of Sciences of the United States of America 91, 7787–7791 (1994).

5 Butterfield, D. A., Reed, T. & Sultana, R. Roles of 3-nitrotyrosine- and 4-hydroxynonenal-modified brain proteins in the progression and pathogenesis of Alzheimer’s disease. Free radical research 45, 59–72, doi:10.3109/10715762.2010.520014 (2011).

6 Greilberger, J. et al. Malondialdehyde, carbonyl proteins and albumin-disulphide as useful oxidative markers in mild cognitive impairment and Alzheimer’s disease. Free radical research 42, 633–638, doi:10.1080/10715760802255764 (2008).

7 Rosborough, K., Patel, N. & Kalia, L. V. alpha-Synuclein and Parkinsonism: Updates and Future Perspectives. Current neurology and neuroscience reports 17, 31, doi:10.1007/s11910-017-0737-y (2017).

8 Sulistio, Y. A. & Heese, K. The Ubiquitin-Proteasome System and Molecular Chaperone Deregulation in Alzheimer’s Disease. Molecular neurobiology 53, 905–931, doi:10.1007/s12035-014-9063-4 (2016).

9 Levine, B. & Kroemer, G. Biological Functions of Autophagy Genes: A Disease Perspective. Cell 176, 11–42, doi:10.1016/j.cell.2018.09.048 (2019).

10 Oury, T. D., Tatro, L., Ghio, A. J. & Piantadosi, C. A. Nitration of tyrosine by hydrogen peroxide and nitrite. Free radical research 23, 537–547 (1995).

11 D’Agostino, D. P., Olson, J. E. & Dean, J. B. Acute hyperoxia increases lipid peroxidation and induces plasma membrane blebbing in human U87 glioblastoma cells. Neuroscience 159, 1011–1022, doi:10.1016/j.neuroscience.2009.01.062 (2009).

12 Ayala, A., Munoz, M. F. & Arguelles, S. Lipid peroxidation: production, metabolism, and signaling mechanisms of malondialdehyde and 4-hydroxy-2-nonenal. Oxidative medicine and cellular longevity 2014, 360438, doi:10.1155/2014/360438 (2014).

13 Gottesman, M. M., Fojo, T. & Bates, S. E. Multidrug resistance in cancer: role of ATP-dependent transporters. Nat Rev Cancer 2, 48–58, doi:10.1038/nrc706 (2002).

14 Safaei, R. et al. Abnormal lysosomal trafficking and enhanced exosomal export of cisplatin in drug-resistant human ovarian carcinoma cells. Molecular cancer therapeutics 4, 1595–1604, doi:10.1158/1535-7163.MCT-05-0102 (2005).

15 Johnstone, R. M. Exosomes biological significance: A concise review. Blood cells, molecules & diseases 36, 315–321, doi:10.1016/j.bcmd.2005.12.001 (2006).

16 Davis, C. H. et al. Transcellular degradation of axonal mitochondria. Proceedings of the National Academy of Sciences of the United States of America 111, 9633–9638, doi:10.1073/pnas.1404651111 (2014).

17 Desir, S. et al. Chemotherapy-Induced Tunneling Nanotubes Mediate Intercellular Drug Efflux in Pancreatic Cancer. Scientific reports 8, 9484, doi:10.1038/s41598-018-27649-x (2018).

18 Melentijevic, I. et al. C. elegans neurons jettison protein aggregates and mitochondria under neurotoxic stress. Nature 542, 367–371, doi:10.1038/nature21362 (2017).

19 Fader, C. M., Sanchez, D., Furlan, M. & Colombo, M. I. Induction of autophagy promotes fusion of multivesicular bodies with autophagic vacuoles in k562 cells. Traffic 9, 230–250, doi:10.1111/j.1600-0854.2007.00677.x (2008).

20 Schonheit, B., Zarski, R. & Ohm, T. G. Spatial and temporal relationships between plaques and tangles in Alzheimer-pathology. Neurobiology of aging 25, 697–711, doi:10.1016/j.neurobiolaging.2003.09.009 (2004).

21 Schwab, C., Steele, J. C. & McGeer, P. L. Pyramidal neuron loss is matched by ghost tangle increase in Guam parkinsonism-dementia hippocampus. Acta neuropathologica 96, 409–416 (1998).

22 Spillantini, M. G., Crowther, R. A., Jakes, R., Hasegawa, M. & Goedert, M. alpha-Synuclein in filamentous inclusions of Lewy bodies from Parkinson’s disease and dementia with lewy bodies. Proceedings of the National Academy of Sciences of the United States of America 95, 6469–6473 (1998).

23 Mijaljica, D., Prescott, M. & Devenish, R. J. The intricacy of nuclear membrane dynamics during nucleophagy. Nucleus 1, 213–223, doi:10.4161/nucl.1.3.11738 (2010).

24 Papandreou, M. E. & Tavernarakis, N. Nucleophagy: from homeostasis to disease. Cell death and differentiation 26, 630–639, doi:10.1038/s41418-018-0266-5 (2019).

25 Brophy, P. M. & Barrett, J. Detoxification of secondary products of lipid peroxidation in the cytosol of a mouse fibroblast cell line. Biochemistry and cell biology = Biochimie et biologie cellulaire 68, 1288–1291 (1990).

26 Rittner, H. L. et al. Aldose reductase functions as a detoxification system for lipid peroxidation products in vasculitis. The Journal of clinical investigation 103, 1007–1013, doi:10.1172/JCI4711 (1999).

27 Mann, S. S. & Hammarback, J. A. Molecular characterization of light chain 3. A microtubule binding subunit of MAP1A and MAP1B. The Journal of biological chemistry 269, 11492–11497 (1994).

28 Xie, R. et al. Microtubule-associated protein 1S (MAP1S) bridges autophagic components with microtubules and mitochondria to affect autophagosomal biogenesis and degradation. The Journal of biological chemistry 286, 10367–10377, doi:10.1074/jbc.M110.206532 (2011).

29 Ponpuak, M., et al. Secretory autophagy. Current opinion in cell biology 35, 106–116, doi:10.1016/j.ceb.2015.04.016 (2015).

30 Holt, O. J., Gallo, F. & Griffiths, G. M. Regulating secretory lysosomes. Journal of biochemistry 140, 7–12, doi:10.1093/jb/mvj126 (2006).

31 Buschow, S. I., Liefhebber, J. M., Wubbolts, R. & Stoorvogel, W. Exosomes contain ubiquitinated proteins. Blood cells, molecules & diseases 35, 398–403, doi:10.1016/j.bcmd.2005.08.005 (2005).

32 Katzenell, S. & Leib, D. A. Herpes Simplex Virus and Interferon Signaling Induce Novel Autophagic Clusters in Sensory Neurons. Journal of virology 90, 4706–4719, doi:10.1128/JVI.02908-15 (2016).

33 Ma, L. et al. Discovery of the migrasome, an organelle mediating release of cytoplasmic contents during cell migration. Cell research 25, 24–38, doi:10.1038/cr.2014.135 (2015).

34 Valcz, G. et al. En bloc release of MVB-like small extracellular vesicle clusters by colorectal carcinoma cells. Journal of extracellular vesicles 8, 1596668, doi:10.1080/20013078.2019.1596668 (2019).

35 Li, S. C. & Kane, P. M. The yeast lysosome-like vacuole: endpoint and crossroads. Biochimica et biophysica acta 1793, 650–663, doi:10.1016/j.bbamcr.2008.08.003 (2009).

36 Roeder, A. D. & Shaw, J. M. Vacuole partitioning during meiotic division in yeast. Genetics 144, 445–458 (1996).

37 Wang, L., et al. alpha-synuclein multimers cluster synaptic vesicles and attenuate recycling. Current biology : CB 24, 2319–2326, doi:10.1016/j.cub.2014.08.027 (2014).

38 Duan, J., Ying, Z., Su, Y., Lin, F. & Deng, Y. alpha-Synuclein binds to cytoplasmic vesicles in U251 glioblastoma cells. Neuroscience letters 642, 148–152, doi:10.1016/j.neulet.2017.01.067 (2017).

39 Dante, S., Hauss, T., Brandt, A. & Dencher, N. A. Membrane fusogenic activity of the Alzheimer’s peptide A beta(1-42) demonstrated by small-angle neutron scattering. Journal of molecular biology 376, 393–404, doi:10.1016/j.jmb.2007.11.076 (2008).

40 Saman, S. et al. Exosome-associated tau is secreted in tauopathy models and is selectively phosphorylated in cerebrospinal fluid in early Alzheimer disease. The Journal of biological chemistry 287, 3842–3849, doi:10.1074/jbc.M111.277061 (2012).

41 Dawson, G. R. et al. Age-related cognitive deficits, impaired long-term potentiation and reduction in synaptic marker density in mice lacking the beta-amyloid precursor protein. Neuroscience 90, 1–13 (1999).

42 Dawson, H. N. et al. Inhibition of neuronal maturation in primary hippocampal neurons from τ deficient mice. Journal of cell science 114, 1179 (2001).

43 Abeliovich, A. et al. Mice lacking alpha-synuclein display functional deficits in the nigrostriatal dopamine system. Neuron 25, 239–252 (2000).

44 Herrup, K. The case for rejecting the amyloid cascade hypothesis. Nature neuroscience 18, 794–799, doi:10.1038/nn.4017 (2015).

45 Meldolesi, J. Exosomes and Ectosomes in Intercellular Communication. Current biology : CB 28, R435–R444, doi:10.1016/j.cub.2018.01.059 (2018).

46 Hargett, L. A. & Bauer, N. N. On the origin of microparticles: From “platelet dust” to mediators of intercellular communication. Pulmonary circulation 3, 329–340, doi:10.4103/2045-8932.114760 (2013).

47 Paluch, E., Piel, M., Prost, J., Bornens, M. & Sykes, C. Cortical actomyosin breakage triggers shape oscillations in cells and cell fragments. Biophysical journal 89, 724–733, doi:10.1529/biophysj.105.060590 (2005).

48 Headley, M. B. et al. Visualization of immediate immune responses to pioneer metastatic cells in the lung. Nature 531, 513–517, doi:10.1038/nature16985 (2016).

49 Patheja, P. & Sahu, K. Macrophage conditioned medium induced cellular network formation in MCF-7 cells through enhanced tunneling nanotube formation and tunneling nanotube mediated release of viable cytoplasmic fragments. Experimental cell research 355, 182–193, doi:10.1016/j.yexcr.2017.04.008 (2017).

50 Goyal, M. S. et al. Loss of Brain Aerobic Glycolysis in Normal Human Aging. Cell metabolism 26, 353–360 e353, doi:10.1016/j.cmet.2017.07.010 (2017).

51 Vlassenko, A. G. et al. Aerobic glycolysis and tau deposition in preclinical Alzheimer’s disease. Neurobiology of aging 67, 95–98, doi:10.1016/j.neurobiolaging.2018.03.014 (2018).

52 Fu, H. L. et al. TET1 exerts its tumor suppressor function by interacting with p53-EZH2 pathway in gastric cancer. Journal of biomedical nanotechnology 10, 1217–1230 (2014).

